# Beyond variance: simple random distributions are not a good proxy for intraspecific variability in systems with environmental structure

**DOI:** 10.1101/2022.08.06.503032

**Authors:** Camille Girard-Tercieux, Ghislain Vieilledent, Adam Clark, James S. Clark, Benoit Courbaud, Claire Fortunel, Georges Kunstler, Raphaël Pélissier, Nadja Rüger, Isabelle Maréchaux

## Abstract

The role of intraspecific variability (IV) in shaping community dynamics and species coexistence has been intensively discussed over the past decade and modelling studies have played an important role in that respect. However, these studies often implicitly assume that IV can be represented by independent random draws around species-specific mean parameters. This major assumption has largely remained undiscussed, although a great part of observed IV is structured in space or time, in particular when environmental dimensions that influence individual performance are imperfectly characterised or unobserved in the field. To test the impact of this strong assumption on the outcome of community dynamics models, we designed a simulation experiment where we varied the level of knowledge of the environment in virtual communities, resulting in different relative importance of explained *vs* unexplained spatial individual variation in performance. We used a community dynamics simulator to generate communities where the unexplained individual variation is, or is not, added as an unstructured random noise. Communities simulated with unstructured IV never reached the community diversity and composition of those where all the variation was explained and structured (perfect knowledge model). This highlights that incorporating unstructured IV (*i.e*. a random noise) to account for unexplained (but structured) variation can lead to incorrect simulations of community dynamics. In addition, the effects of unstructured IV on community diversity and composition depended on the relative importance of structured *vs* unstructured IV, *i.e*. on the level of knowledge of the environment, which may partly explain the contrasting results of previous studies on the effect of IV on species coexistence. In particular, the effect of unstructured IV on community diversity was positive when the proportion of structured IV *vs* unstructured IV in the model was low, but negative when this proportion was high. This is because unstructured random noise can either limit the competitive exclusion of inferior competitors in low dimensions or destabilise tight niche partitioning in high dimension. Our study suggests that it is crucial to account for the sources and structure of observed IV in real communities to better understand its effect on community assembly and properly include it in community dynamics models.

## Introduction

The role of intraspecific variability (IV) in shaping community dynamics has been intensively discussed over the past decade (Bolnick et al. 2011; Albert et al. 2011; Violle et al. 2012; Des Roches et al. 2018; Raffard et al. 2019). Observed IV, *i.e*. the variability among measured individual attributes (functional or demographic traits, or any proxy of individual performance) within a species has indeed been reported to be large within communities (Siefert et al. 2015; Poorter et al. 2018). Modelling studies have played an important role to decipher the effect of IV on species coexistence (*e.g*. Lichstein et al. 2007; Vieilledent et al. 2010; Courbaud et al. 2012; Hart et al. 2016; Uriarte and Menge 2018; Crawford et al. 2019), offering opportunities of virtual experiments out of the scope of empirical approaches. These studies have led to contrasting results however, letting the debate unresolved: IV could either (i) blur species differences, thus promoting transient or unstable coexistence (Vieilledent et al. 2010; Crawford et al. 2019), (ii) disproportionately advantage the strongest competitor, thus hindering coexistence (Courbaud et al. 2012; Hart et al. 2016), or (iii) promote coexistence in specific spatial configurations (Uriarte and Menge 2018). While a unifying framework differentiating whether IV affects niche traits or hierarchical traits has been recently proposed to explain these discrepancies (Stump et al. 2022), a major assumption usually made in modelling studies, namely that IV is unstructured in space or time and can be represented by independent random draws around species-specific mean parameters, remains largely undiscussed (Girard-Tercieux et al. 2023).

The IV observed in individual attributes is not necessarily purely random and can emerge from various genetic and environmental processes (Violle et al. 2012; Moran et al. 2016). Most of these additional processes are unlikely to generate unstructured IV in the form of a random noise, whereby the site and date of measurement would have no influence on the measured attribute value (unstructured IV, henceforth denoted uIV). Previous works have already explored the role of genetically heritable traits variability (Ehlers et al. 2016). In contrast, much less attention has been given to IV generated by structured variation of environmental gradients in space or time (structured IV, henceforth denoted sIV). It is, however, well-known that many species attributes respond to environmental gradients (Bonnier 1890; Kropotkine 2015; Jung et al. 2010; Niinemets 2015; Rixen et al. 2022). As a result, high-dimensional (and potentially unobserved) variation of the environment can lead to large observed IV. For instance, in a highly controlled clonal experiment, IV in tree growth within clones was larger than genetically-driven IV between clones (Girard-Tercieux et al. 2023). Indeed, differences in attributes among conspecific individuals can result from differences in environmental dimensions that are unobserved or mischaracterised due to a mismatch between the individual scale and the scale of the measurements. Consequently, these observed differences do not necessarily mean that conspecific individuals substantially differ in their response to the environment. While it is widely accepted that environmentally-driven sIV is ubiquitous in natural communities (Nicotra et al. 2010), the consequences of its substitution by random uIV on species coexistence and community dynamics remain to be thoroughly tested in models (Clark 2010; Girard-Tercieux et al. 2023).

Here, we explore the effect of considering IV either as structured by environmental dimensions (sIV) or as an unstructured random noise (uIV), through a virtual experiment designed to provide a first proof-of-concept, performed using a simulator of community dynamics. To do so, we first created a virtual plant community, where individual performance is fully determined by specieslevel responses to 15 environmental dimensions varying in space (Fig. 1A). This extreme scenario, although unrealistic regarding its level of environmental determinism, was subsequently used as a reference (henceforth denoted *Perfect knowledge model*) in our virtual experiment. We then considered imperfect knowledge models, where this 15-dimensional individual performance is estimated using 0 to 15 supposedly “observed” environmental dimensions, while the remaining IV (or unexplained variation) resulting from the effect of “unobserved” environmental dimensions, is ignored (*Imperfect knowledge models without uIV*) or is included as random unstructured IV (*Imperfect knowledge models with uIV*, Fig. 1B). These three performance models are used to independently run the same community dynamics simulator in order to compare their effects on species coexistence and community dynamics.

**Figure 1:**
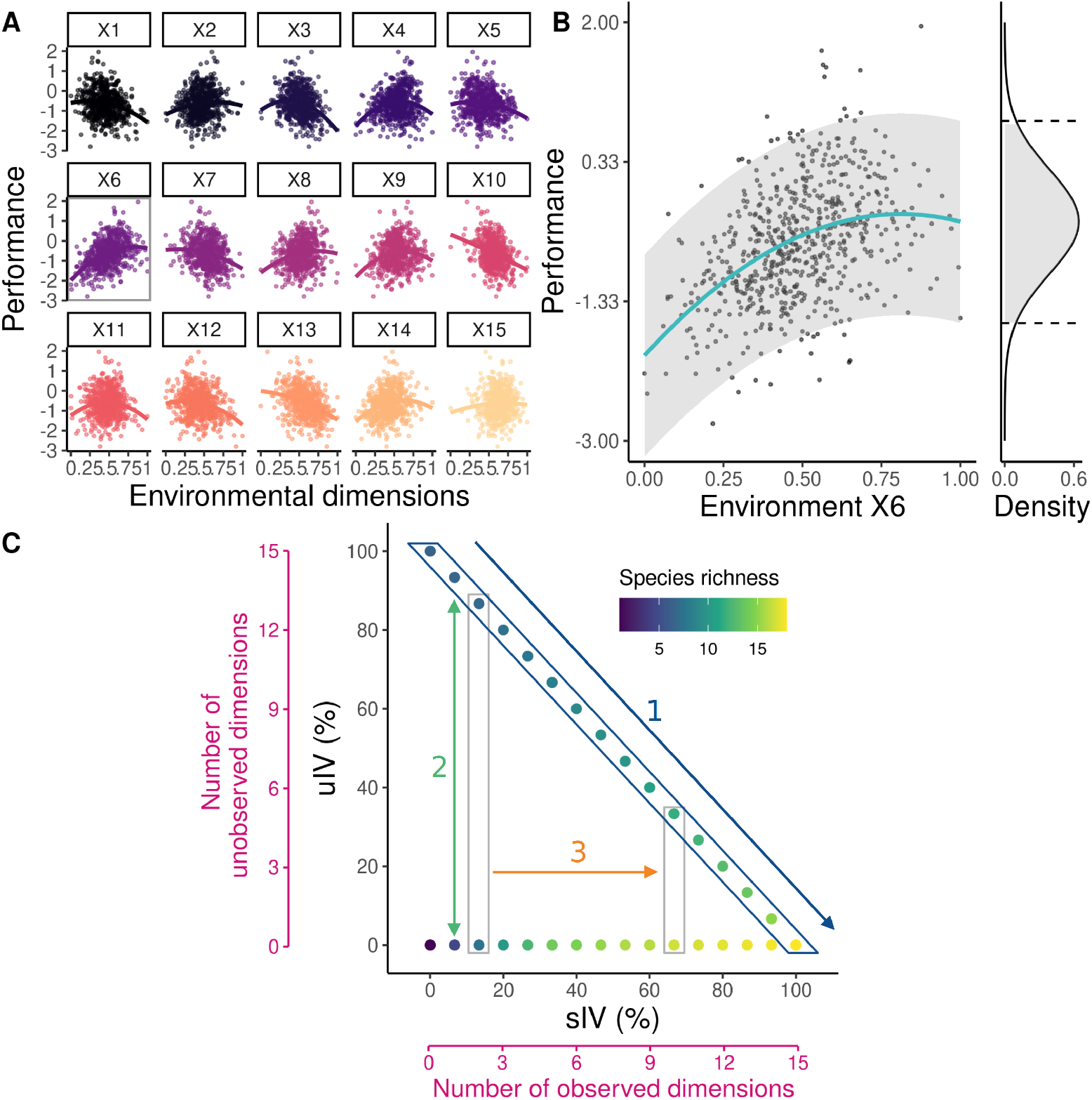
Conceptual framework. Consider an environment that is varying spatially in many dimensions, X1 to X15. Each dimension influences individual performance in a species-specific way, as illustrated in A for one species (where the variation of performance with all environmental variables is projected separately for each variable in 2-dimensional plots). In practice, several of these environmental dimensions are often unobserved in the field. The effect of these unobserved environmental dimensions on individual performance results in an observed intraspecific variability (IV) in the species response to observed dimensions. As an illustration, in B, only X6 is observed and used to fit a polynomial function to the performance data (teal curve), and the remaining variation is estimated through a variance term (gray envelope). This variability is often represented as a probability distribution, which is used to simulate the variation in performance among conspecific individuals through random draws that are independent and unstructured in space (density panel in B). We propose a framework to assess the consequences of representing the variation resulting from unobserved environmental dimensions, which is structured in space and time, by such unstructured IV (uIV) on community dynamics, and how these consequences vary with the level of knowledge of the environment (C). To do so, we varied from 0 to 15 the number of dimensions that are observed and used for estimating the 15-dimensional performance (panel B providing an example with one dimension). By increasing the number of observed dimensions, we thus increased the proportion of structured IV (sIV) that is accounted for in estimating individual performance (C, horizontal axis; see also Fig. 2). For a given number of observed dimensions, or % of sIV, the variation resulting from unobserved dimensions can be either added as uIV or not (C, vertical axis). For each way to estimate performance (with uIV or not, and with different numbers of observed dimensions), we simulated community dynamics using the same simulator. We then compared the simulated communities in terms of diversity and composition (*e.g*. species richness in colored points in C). By comparing communities simulated with uIV with the one with 100 % sIV (arrow 1) we tested the effect of substituting sIV with uIV on community dynamics. By comparing communities simulated with and without uIV, for a given % of sIV (arrow 2), we mimicked the approach of previous modelling studies testing the effect of intraspecific variability on community dynamics and species coexistence. By comparing this difference between communities simulated with and without uIV across different % of sIV (arrow 3), we tested whether the results of previous studies can be influenced by the level of knowledge of the environment.

Specifically, we are asking two questions. First, how well does random unstructured IV (uIV) mimic the effect of environmentally-driven structured IV (sIV) on diversity and community composition? To answer this question, we compare communities simulated under the *Perfect knowledge model* and under *Imperfect knowledge models with uIV*. Importantly, these models share the same amount of total variation across individuals, but partitioned differently between sIV and uIV, depending on the amount of knowledge of the environment, *i.e*. on the number of “observed” environmental dimensions (Fig. 1C, arrow 1). Second, how does the effect of adding uIV on diversity and community composition vary with the knowledge of the environment (Fig. 1C, arrow 2), *i.e*. with the relative importance of sIV and uIV in our model? To answer this question, we compare pairs of models with the same amount of sIV, *i.e*. with the same knowledge of the environment, but including or excluding uIV. This latter comparison corresponds to the approaches proposed in previous studies testing the effects of IV on coexistence (Vieilledent et al. 2010; Courbaud et al. 2012; Hart et al. 2016).

## Materials and methods

### Environmental variables

We considered a grid of *M* = 25 × 25 = 625 sites. Each site *m* was characterised by *K* = 15 environmental variables *x*_1_, …, *x*_*K*_. To confer some realism to our virtual experiment and the resulting illustrations (Fig. S5.17), each environmental variable was spatially auto-correlated, as it is often the case in nature (Tymen et al. 2017; Zellweger et al. 2019), and independently derived from a conditional autoregressive model, with a normal distribution centered on 0 and of variance 1. Therefore, environmental variables were not uniformly distributed, some habitats being more frequent than others. Environmental variables were then rescaled to [0, 1] to ensure that each variable had the same effect on species performance on average. We here assumed that the environmental variables do not vary in time and therefore restricted our experiment to environmental variation in space.

### Individual performance

For parsimony’s sake, we here focus on an attribute (or trait) that has a direct link with performance, hereafter “individual performance”. We considered *J* = 20 species, whose individual performances were computed in three alternative ways, as follows.

### The *Perfect knowledge model*

We first considered a simple model representing the functioning of a plant community in a hypothetical world where all determinants of individual performance would be environmental and known named *Perfect knowledge model* and henceforth considered as the reference. Individuals within a species did not have any intrinsic differences and could therefore be considered as clones, and we assumed no genetic variation among individuals. We considered that the environment was multidimensional and partitioned among species. To this end, in this model, the performance of an individual *i* of species *j* (*j* ∈ [1, …, *J*]) was maximal at one point in the multidimensional environmental space, denoted 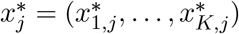. For an environmental axis *k* (*k* ∈ [1, …, *K*]), 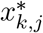 was drawn in a uniform distribution in [0, 1]. Then, the performance of an individual *i* of species *j* on site *m, p*_*i,j,m*_, was computed as the opposite of the normalised Euclidean distance between 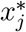 and the local environment at the site where the individual resided, *x*_*m*_ = (*x*_1,*m*_, …, *x*_*K,m*_) (Eq.1).

Therefore, at each site, one species outperformed all the others. The number of sites where each species had the highest performance varied between species, since the environmental variables were not uniformly distributed. For some species, there was no site where they were the most competitive. Importantly, all individuals of a given species *j* responded in the same way to the environment, the performance of conspecifics differing only because they resided in a different environment. Individual variation was thus fully environmentally-driven and structured in space (0% uIV and 100% sIV in Fig. 1C).

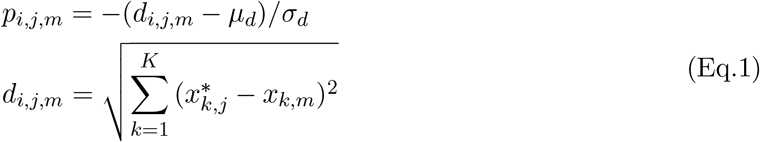

where *µ*_*d*_ and *σ*_*d*_ are the mean and variance of *d*_*j,m*_ across all species *j* and sites *m*.

### The *Imperfect knowledge models*

As it is typically unfeasible to fully characterise all relevant dimensions of the environment at fine scales in the field, we then assumed that only *n*_obs_ *<* 15 environmental variables were measured and accounted for when estimating individual performance. These performances were thus estimated from a statistical model fitting the individual performance *p*_*i,j,m*_ provided by the *Perfect knowledge model* (Eq.1, representing what actually happens in the field and is measured, assuming no measurement error) against the *n*_obs_ observed environmental variables (Fig. 1). We considered the common case where ecologists, in absence of exact knowledge of the underlying processes, here depicted by the *Perfect knowledge model*, assume a quadratic relationship between performance and each observed environmental variable, thus approaching the triangular shape (*i.e*. increasing then decreasing piecewise linear) of the actual relationship of the *Perfect knowledge model* (Eq.2). The use of an intercept to estimate species performance in averaged environmental conditions is also a common practice among ecologists.

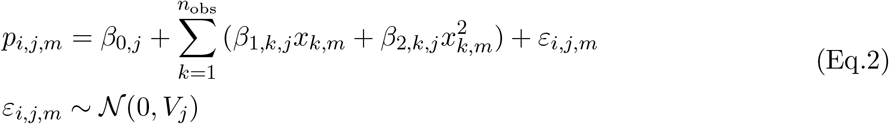

This statistical model was fitted using the “lm” function of the “stats” R package. Species parameters (*β*_*j*_ = *{β*_0,*j*_, *β*_1,*k,j*_, *β*_2,*k,j*_}) and residuals *ε*_*i,j,m*_ were retrieved. In this model, *V*_*j*_ represented an unstructured observed IV which was estimated for each species *j*. This variability emerged from the spatial variation in environmental variables that were not measured and accounted for, namely 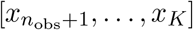.

In *Imperfect knowledge models*, the individual performance 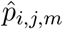 was computed with the parameters obtained at different levels of knowledge of the environment using Eq.2, *i.e*. with *n*_obs_ varying from 0 to 15. *ε*_*i,j,m*_ thus accounted for the *K* − *n*_obs_ unobserved environmental variables, respectively. In the *Imperfect models without uIV*, the residual variation, *ε*_*i,j,m*_, was neglected (Eq.3), while in the *Imperfect knowledge models with uIV*, it was included as a random noise *δ*_*i,j,m*_ generated through independent individual draws in a normal distribution of variance 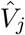 (Eq.4). The *Imperfect knowledge models with uIV* therefore shares the same amount of total variation across individuals with the *Perfect knowledge model*, but partitioned differently between sIV and uIV: for a given number of observed environmental dimensions, random IV *V*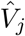 was used as a substitute of the environmental variation that was not observed.

Importantly, in the *Imperfect knowledge models without uIV*, conspecific individuals responded similarly to the environment as in the *Perfect knowledge model* for the observed environmental dimensions, but lacking information on the other environmental dimensions (0% uIV in Fig. 1C).

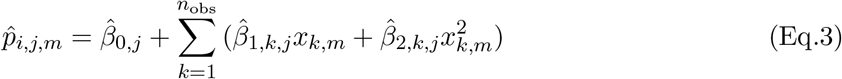

In contrast to the *Perfect knowledge model* and the *Imperfect knowledge models without uIV*, in the *Imperfect knowledge models with uIV*, conspecific individuals could perform differently in the same environment (0 to 100% uIV in Fig. 1C), due to the contribution of the random term *δ*_*i,j,m*_ to the performance. Note that this contribution, which mimics what was typically done in previous studies, can lead to a trend of increasing average performance over iterations in simulations of community dynamics, especially due to the unbounded, normal distribution of *δ*_*i,j,m*_.

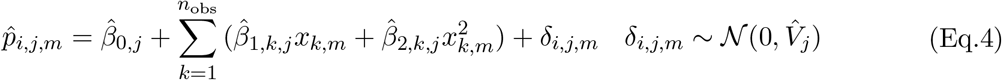

The three types of performance models (Eq.1, Eq.3, Eq.4) were then implemented in the same simulator of community dynamics, in order to disentangle the effects of random, unstructured IV on the one hand, and of the imperfect characterisation of the environment on community dynamics and species coexistence on the other hand. Note that, when comparing simulation outcomes with performance computed with the Perfect Knowledge and the Imperfect Knowledge models, differences can actually result from two main aspects: (i) the model mis-specification (*i.e*. the use of a quadratic function including an intercept, Eq.2, Eq.3, Eq.4, instead of a distance, Eq.1), and (ii) the number of observed dimensions. Change in the latter actually results in a change in the proportion of uIV, but also in the number of variables that can be used to compute the site-dependent (or environment-dependent) part of performance. Moreover, the presence of an intercept in models (Eq.2, Eq.3, Eq.4) can lead to a hierarchy in species performances fostering species extinctions. We tested the magnitude of the effect of the intercept and of the model mis-specification, using an Imperfect knowledge model with a distance function, with and without intercepts, which did not substantially change our results (see Reviews and discussion below).

### Community dynamics simulation

Our simulator of community dynamics was inspired by Hurtt and Pacala (1995). However, several of our modelling choices differed. First, we explicitly used several environmental dimensions to account for niche multidimensionality, while they used a one-dimensional environmental index. Second, we randomly drew species optima, therefore leading to various sizes of the environmental space where each species outperforms all the others, while they used equally wide ecological niches across species. This allowed us to test several configurations of niche partitioning. Finally, mortality and recruitment were stochastic in their model, while we chose a deterministic process to stabilise coexistence and limit the sources of uncertainty to the effect of IV, although we also tested a stochastic alternative (see details below).

For a given simulation of community dynamics, the simulated community was initialised with ten individuals of each of the 20 species, located randomly in the landscape. The performance of these individuals was computed using either the *Perfect knowledge model* (Eq.1), an *Imperfect knowledge model without uIV* (Eq.3), or an *Imperfect knowledge model with uIV* (Eq.4). Mortality events resulted in vacant sites for which species then competed for recruitment. To test the robustness of our results to the choices made in building the community dynamics simulator, we implemented alternative ways to simulate mortality and fecundity. In the following formulas, performance *p*_*i,j,m*_ is replaced by 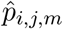 for *Imperfect knowledge models*. For mortality, we explored the three following approaches: (i) the one percent less performing individuals in the landscape die at each timestep, henceforth denoted *deterministic mortality* ; (ii) one percent of the individuals die at each timestep, and the probability *θ*_*i,j,m*_ of each individual *j* to die is inversely proportional to its performance, *θ*_*i,j,m*_ = logit^−1^(0.5 × *p*_*i,j,m*_), henceforth denoted *stochastic mortality* ; (iii) *θ*_*i,j,m*_ is computed as a function of individual performance, *θ*_*i,j,m*_ = logit^−1^(logit(0.01) − 0.5 × *p*_*i,j,m*_), henceforth denoted *logistic stochastic mortality*. Death events are then drawn in a binomial distribution *B*(*n*_*s*_, *θ*) with *θ* the vector of all *θ*_*i,j,m*_. For fecundity, we explored the two following approaches: (i) the number of propagules *λ*_*j,t*_ depends on species abundance *A*_*j,t*_: *λ*_*j,t*_ = round(0.5 × *A*_*j,t*_), henceforth denoted the *abundance-dependent fecundity* ; or (ii) each species present in the community produces ten offspring per timestep, henceforth denoted the *fixed fecundity*. In both cases, propagules were then randomly distributed among all vacant sites. If several propagules landed on the same vacant site, the propagule with the highest individual performance outcompeted the others and won the site. A species that was not the best at a site could win “by forfeit” and be recruited at this site. When individual performance was computed using the *Perfect knowledge model*, the colonisation of a vacant site only depended on the species optima. When individual performance was computed using an *Imperfect knowledge model without uIV*, this colonisation depended on the estimated species parameters (the 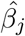), and, for an *Imperfect knowledge model with uIV*, also on a random individual variation (the random term *δ*_*i,j,m*_), that enabled potential inversions of competitive hierarchy locally. Overall, multidimensional niche partitioning and environmental filtering were the main coexistence mechanisms within the simulated communities: mortality and recruitment were controlled by performance, which depended on the local environment in a species-specific way. Therefore, individuals that were maintained and recruited on a site were filtered by the environment, and performance on each site increased rapidly. As population sizes were relatively low, communities were also subject to ecological drift (*i.e*. extinctions due to demographic stochasticity). Note that our community dynamics simulator is spatially-implicit, *i.e*. the fate of an individual on a site does not depend on its neighbours neither on the environment in the neighbourhood (the spatial autocorrelation of the environmental variables does not directly influence in the dynamics and here was only used for illustration, Fig. S5.17). Spatial processes that could contribute to species coexistence (*e.g*. Wiegand et al. 2021*) were thus absent from our simulator, whose aim was not to provide all potential coexistence mechanisms. When using the performance models without uIV (i.e. Perfect knowledge* and *Imperfect knowledge models without uIV*), each species tends to occupy its preferred habitat defined by its optima (perfectly or imperfectly estimated) in many environmental dimensions. It should be noted that species favorable habitats were not equally frequent across species, thus intrinsically defining rare and dominant species in the landscape. Few species that had a rare favorable habitat and whose initial individuals randomly landed on unfavorable sites, could be excluded from the community.

As most results remained qualitatively unchanged across the different alternatives for simulating mortality and fecundity, we present below the results for the *deterministic mortality* and the *abundance-dependent fecundity* only, and refer the reader to Appendix 1 for the other alternatives.

### Experimental setup and analyses

For each model of individual performance and number of observed environmental dimensions, we used ten different Environment × Species optima (*E* × *S*) configurations, each prescribed randomly. Within each *E* × *S* configurations, ten simulations differing only in their initial conditions (location of the initial individuals) were run. Each simulation of community dynamics was run for 10,000 generations (Table 1). The ten *E* × *S* configurations were the same across models of individual performance and number of observed environmental dimensions and the ten initial conditions were the same across *E* × *S* configurations. In total, this led to 3,300 simulations.

**Table 1:**
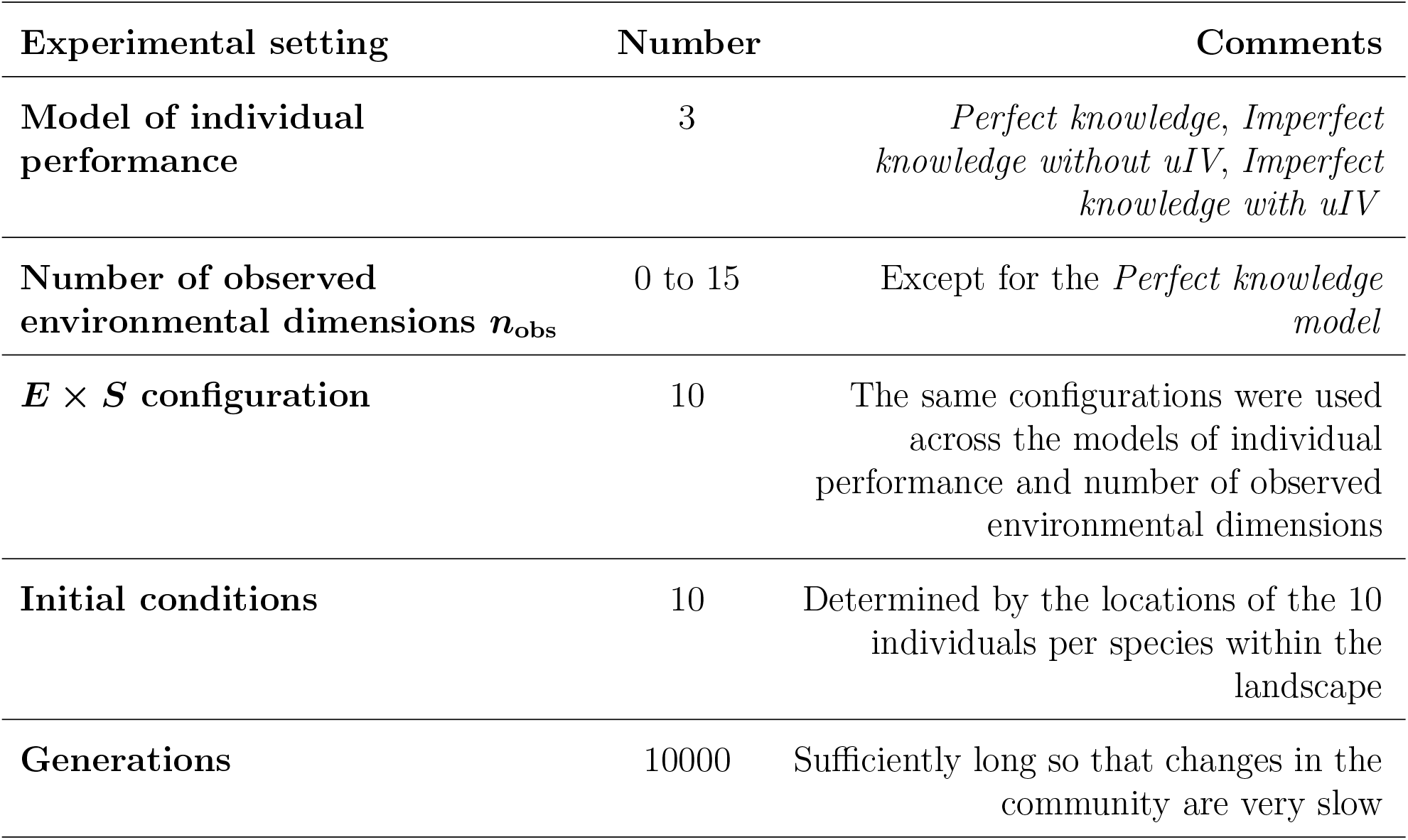
Experimental setup.

In order to compare simulation outputs, we studied several aspects of final communities: (1) community diversity, (2) the similarity in community composition between simulations, and (3) site sorting. Community diversity was estimated using species richness and the Hill-Shannon diversity index (Roswell et al. 2021). Similarity in community composition was estimated as the pairwise percentage similarity of final species abundances between pairs of simulations. For two vectors of species abundances *A* = (*a*_1_, …, *a*_*j*_, …, *a*_*J*_) and *B* = (*b*_1_, …, *b*_*j*_, …, *b*_*J*_), the percentage similarity was computed as

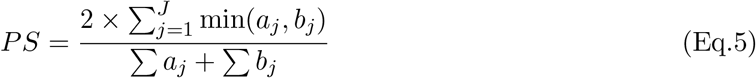

To quantify site sorting, we computed for each simulation the final community mean performance as the performance obtained with the *Perfect knowledge model*, averaged across all individuals at the end of the simulation. This community mean performance thus corresponded to the strength of the environmental filtering in community assembly, *i.e*. the site sorting: the higher the mean performance, the stronger the effect of the environment on community assembly.

## Results

### Final community diversity

Final community diversity, both in terms of species richness and Hill-Shannon index, was lower with unstructured IV than with the *Perfect knowledge model* whatever the number of observed environmental dimensions, *i.e*. whatever the relative importance of structured *vs* unstructured IV (Fig. 3A and B). This diversity increased with the number of observed dimensions. In most cases, adding unstructured IV reduced the community diversity with respect to the corresponding *Imperfect knowledge model without uIV* (Fig. 3C and D). However, this effect varied with the number of observed dimensions (but see in case of alternative mortality implementation, Appendix 1): below 50% of explained variance (*i.e*. up to three observed environmental dimensions, Fig. 2), adding unstructured IV resulted in a higher or similar diversity than with the *Imperfect knowledge models without uIV* (but see in case of alternative mortality implementation, Appendix 1). This difference first decreased and then increased as the number of observed dimensions increased, while staying negative from 3 to 15 observed dimensions.

**Figure 2:**
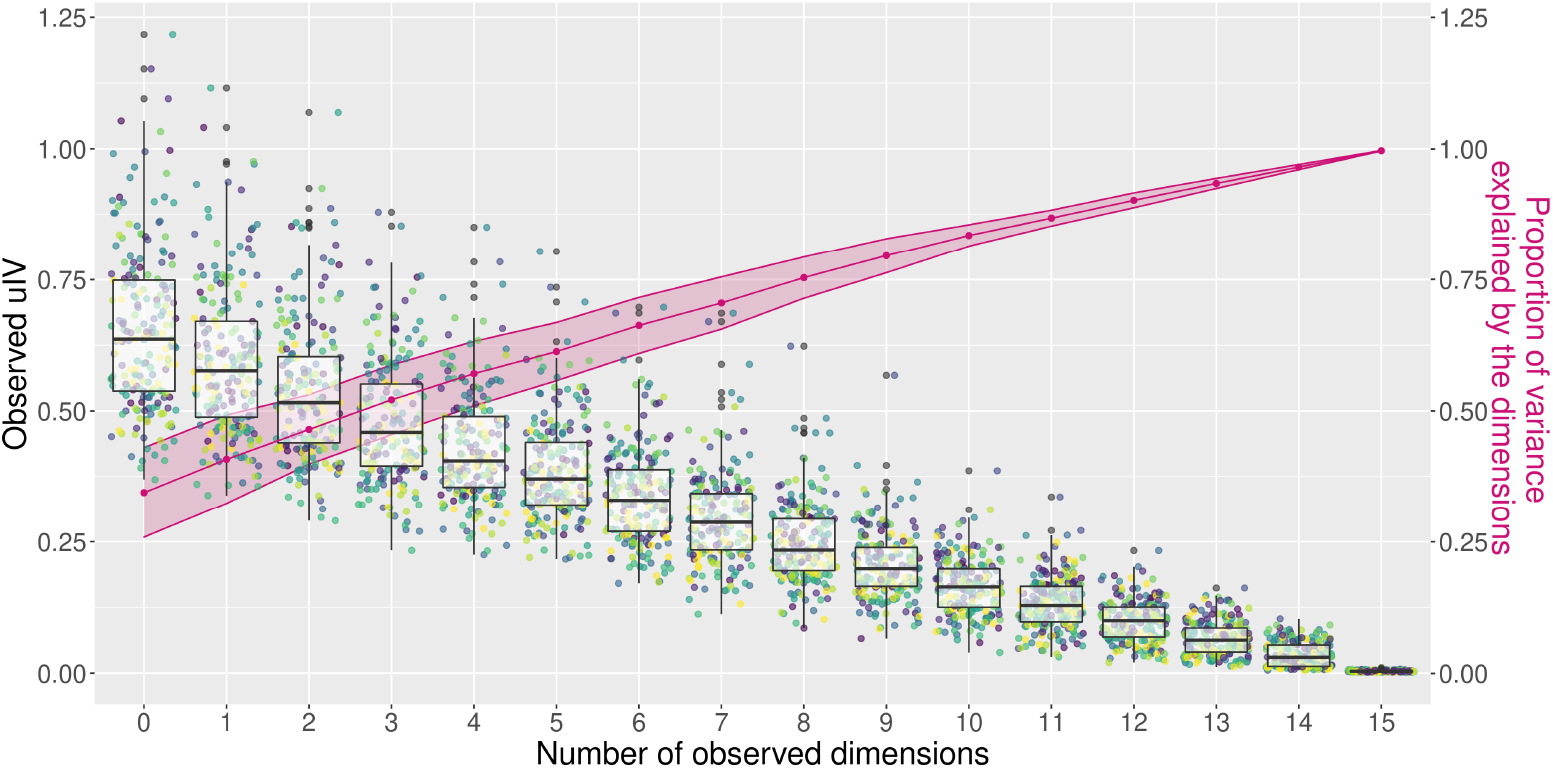
Observed IV depending on the level of knowledge of the environment. Each point represents the unstructured IV inferred for one species, and each colour represents an *E* × *S* configuration (twenty points per colour for the twenty species). Unstructured IV was inferred using a statistical model (Eq.2) taking 0 to 15, out of 15, dimensions into account to fit the performance provided by the *Perfect knowledge model* ; the pink points, curve and ribbon correspond to the mean and standard deviation of the R2 of these statistical models (computed over the ten different configurations for each number of observed dimensions). As expected, observed unstructured IV decreased with the number of observed dimensions, *i.e*. with the level of knowledge of the environment.

**Figure 3:**
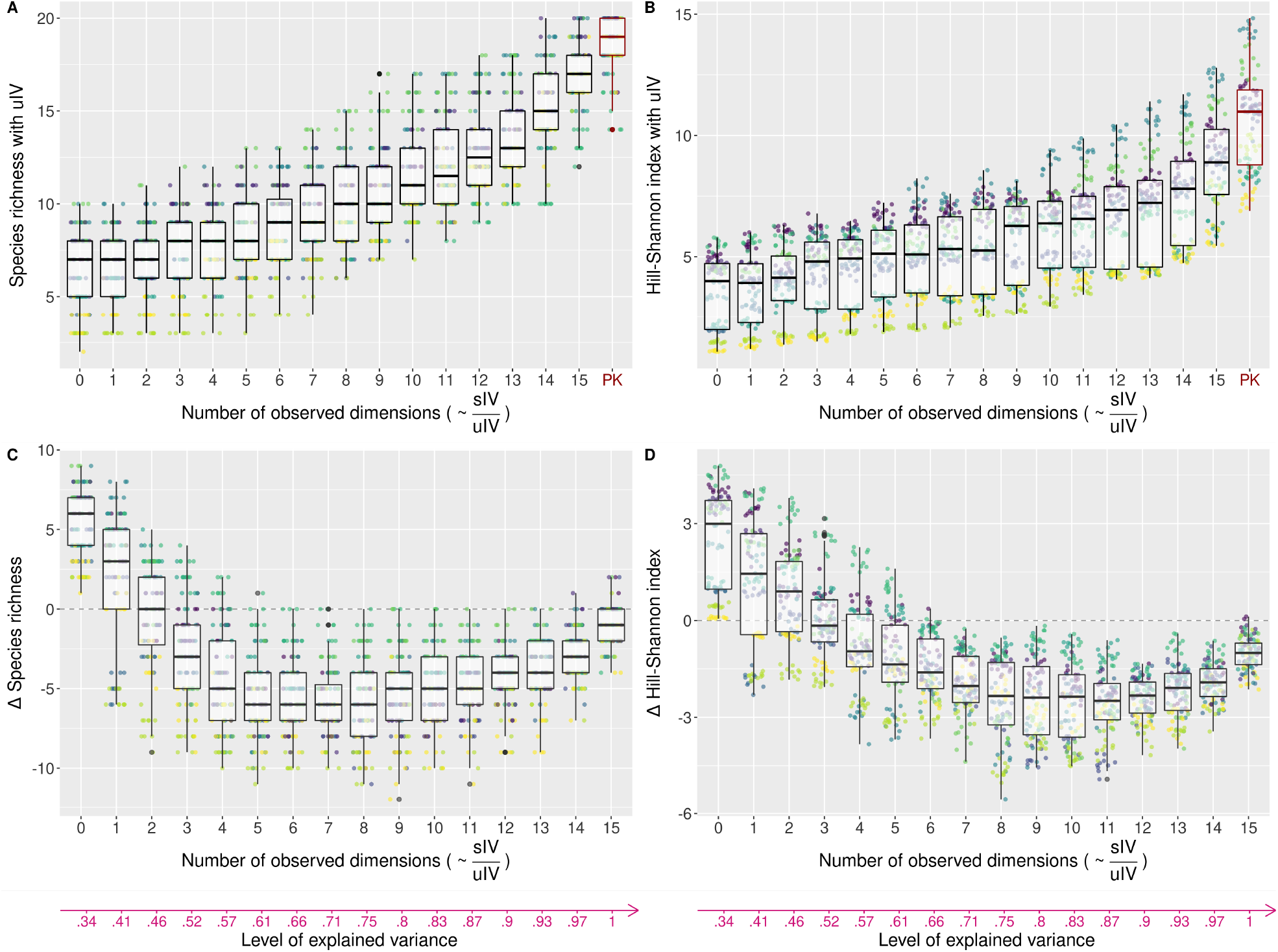
Effect of the structure of individual variation on community diversity. Each point represents the diversity, either computed as the species richness – left panels – or the Hill-Shannon diversity index – right panels – of a final simulated community. Each colour represents an *E* ×*S* configuration (ten points per color, for the ten initial conditions). The horizontal axis corresponds to the number of observed environmental dimensions, which is proportional to the ratio of structured and unstructured IV in the performance models. Each number of observed dimensions corresponds to a level of explained variance in individual performance (see Fig. 2) depicted with the pink arrow at the bottom. The top panels show the final community diversity obtained with the *Imperfect knowledge models with uIV* (0 to 15 observed dimensions) and with the *Perfect knowledge model* (PK, red, far right). This is useful to examine our first question (Fig. 1C, arrow 1). The bottom panels show the difference in the final community diversity obtained with the *Imperfect knowledge models with* and *without uIV*. Points that are above zero (horizontal dashed line) correspond to a higher diversity when adding unstructured IV. This is useful to examine our second question (Fig. 1C, arrows 2 and 3), by comparing the effect of adding unstructured IV at different levels of knowledge of the environment. The *Imperfect knowledge models with uIV* never reached the diversity obtained with the *Perfect knowledge model* (A and B). Moreover, adding unstructured IV as a random noise had an effect on community diversity that varied with the number of observed environmental dimensions (C and D). Results shown here were obtained with a *deterministic mortality* and an *abundance-dependent fecundity* (see main text).

### Final community composition

Similarity (as measured by PS, Eq.5) of the *Imperfect knowledge models with uIV* with the *Perfect knowledge model* was low when few environmental dimensions were observed, *i.e*. when the relative importance of structured (*vs* unstructured) IV was low. This similarity increased with the number of observed dimensions (from 0.55 to 0.9, Fig. 4A). Adding unstructured IV increased the similarity with the *Perfect knowledge model* at low numbers of observed dimensions (from 0 to 2 dimensions, *i.e*. below 50% explained variance) but decreased it at higher numbers of observed dimensions, with respect to the corresponding *Imperfect knowledge model without uIV* (Fig. 4C, but see in case of alternative mortality implementation, Appendix 1). This negative effect became stronger (from 3 to 8 observed dimensions) before becoming weaker (from 9 to 15 observed dimensions). See Appendix 2 for the similarity within models.

**Figure 4:**
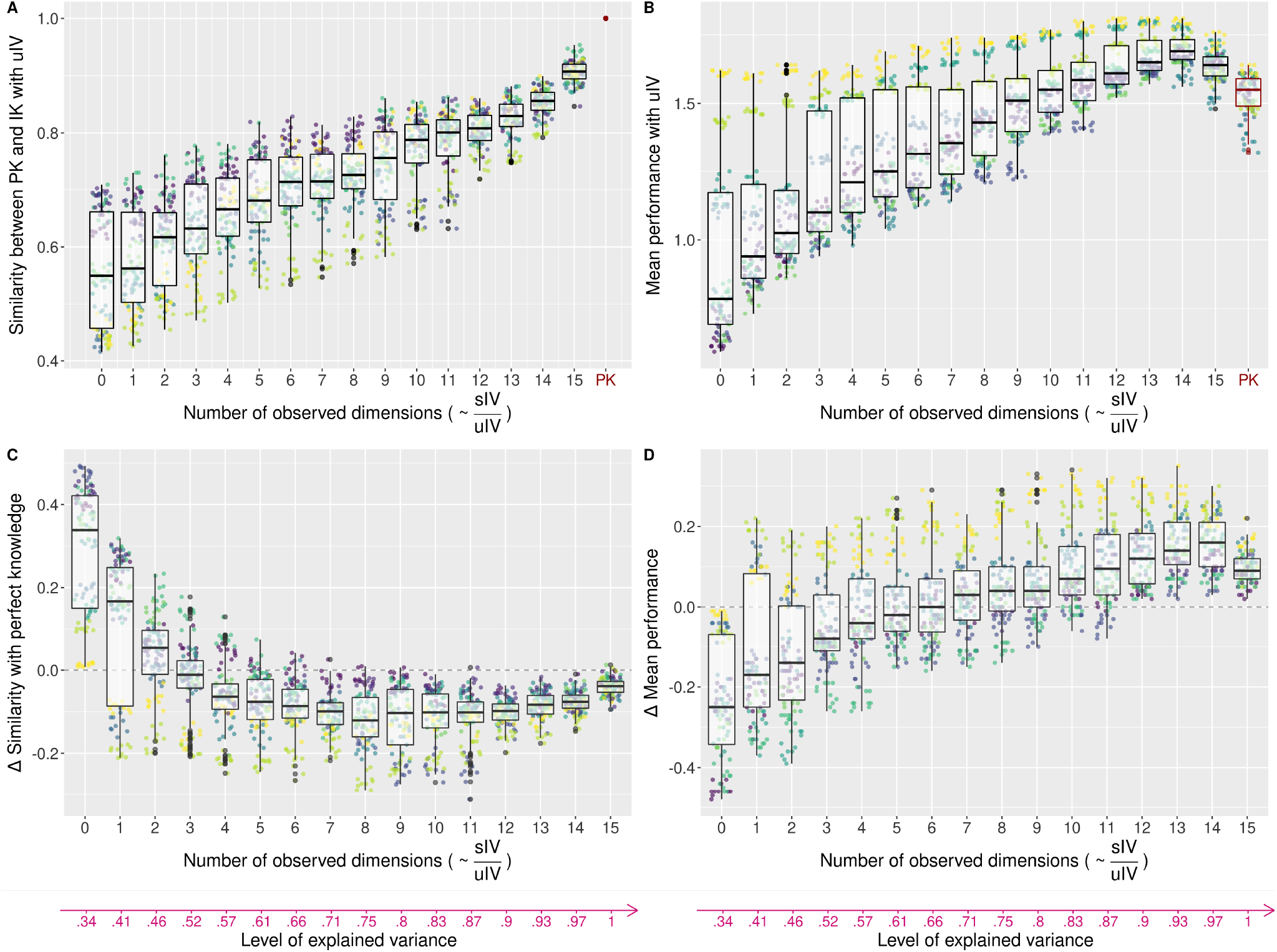
Effect of individual variation on the similarity in final species abundances between models and on the site sorting. Each colour represents an *E* ×*S* configuration. For the similarity left panels -, each point represents the pairwise percentage similarity (PS) in the final species abundances between two simulations with the same *E*× *S* configuration and the same initial conditions (ten points per color), but obtained using the *Perfect knowledge model* on the one hand and one of the *Imperfect knowledge models* on the other hand. For the site sorting right panels -, each point represents the community mean performance of the final communities. This mean performance was calculated with the *Perfect knowledge model* and averaged across all individuals at the end of the simulation. The top panels show these two metrics for communities simulated with the *Imperfect knowledge models with uIV* (0 to 15 observed dimensions) and with the *Perfect knowledge model* (PK, red, far right). The bottom panels show the difference in these metrics for communities obtained with the *Imperfect knowledge models with* and *without unstructured IV*. Points that are above zero (horizontal dashed line) correspond to a higher similarity or mean performance when adding unstructured IV, respectively. The similarity between the *Perfect knowledge model* and the *Imperfect knowledge models with uIV* was low with few observed dimensions and increased with the number of observed dimensions (A). The effect of adding unstructured IV to *Imperfect knowledge models* on the similarity with the *Perfect knowledge model* varied with the number of observed environmental dimensions (C). The mean performance obtained for communities simulated with the *Imperfect knowledge models with uIV* as well as its difference with the *Imperfect knowledge models without uIV* varied with the number of observed dimensions (B, D). Results shown here were obtained with a *deterministic mortality* and an *abundance-dependent fecundity* (see main text).

The mean performance of communities simulated with the *Imperfect knowledge models with uIV* increased with the number of observed dimensions (except between 14 and 15 observed dimensions), *i.e*. with the relative importance of structured and unstructured IV (Fig. 4B). Below ten observed dimensions, it remained lower than that of the communities simulated with the *Perfect knowledge model*, but was higher above ten observed dimensions. Adding unstructured IV decreased the mean performance of the final species community from zero to six observed dimensions but increased it at higher numbers of observed dimensions, with respect to the corresponding *Imperfect knowledge model without uIV* (Fig. 4D). This difference increased with the number of observed dimensions, except between 14 and 15 observed dimensions.

## Discussion

### Random unstructured individual variability generates communities with lower diversity that are dissimilar from communities generated with structured individual variability

Ecologists often have only access to an imperfect characterisation of all the environmental dimensions that actually lead to individual variation, be it due to some overlooked dimensions or variables, or a monitoring at a scale coarser than the one of the variation that actually influences individuals. This mischaracterisation can result in an observed but unexplained intraspecific variability in data. To account for it in community dynamics models, it has often been (implicitly) assumed that some unstructured variation could be added to the explained part of variation to reach the actual observed total variation. To test this assumption, in our simulation experiment, we varied the level of knowledge of the environment and incorporated the remaining (unexplained) variability in individual performance as unstructured noise, thus varying the ratio of structured and unstructured IV. Using a modelling experiment, we showed that this difference in the nature of IV can have strong consequences on community structure and composition.

Within our modelling framework, and compared to the reference communities simulated with a 15-dimensional individual performance, the communities simulated with a performance estimated with fewer dimensions and to which the remaining variance was added as a random noise were less diverse (see also Appendix 3 for further explanation on simulated species richness). Beyond the community diversity *per se*, community composition was dissimilar from the reference when the number of observed dimensions was low, *i.e*. when the relative importance of structured *vs* unstructured IV was low: the strength of environmental filtering in shaping community assembly was too low to generate species abundances similar to the one of the reference communities. As the relative importance of structured IV increased, both the strength of environmental filtering and the similarity of the final species abundances with the reference ones increased.

While the intercepts included in the Imperfect knowledge models resulted in a species hierarchy leading to a drop in species richness at low numbers of observed dimensions, this effect quickly weakened as the number of observed dimensions increased. Overall this effect alone did not explain the observed patterns and our results were qualitatively robust to the use of a distance rather than a quadratic function in Imperfect knowledge models (see Reviews).

Finally, random intraspecific variability is not a good substitute for species response to unobserved environmental dimensions for studying community dynamics. Moreover, interpreting observed IV as unstructured differences in conspecifics’ response to the environment can lead to misinterpretations regarding the ecological mechanisms driving the community dynamics. It would mistake the response of species to environmental variation (a niche mechanism) with random variability (typically a neutral mechanism, *i.e*. affecting all species in the same way), and present IV as a coexistence mechanism *per se* without taking into account the species-specific responses to environmental variations in high dimensions from which IV can actually result. Hence, maintaining the variance observed among individuals is not sufficient to capture the community dynamics, the structure and nature of this individual variability is also critical.

### The effect of adding a random IV depends on the relative importance of structured *vs* unstructured individual variability

Previous modelling studies that explored the role of IV on community dynamics usually did not maintain the total level of variance among individuals. They typically compared communities with and without additional random variability, for the same level of explained individual variation (*Imperfect knowledge model without uIV vs Imperfect knowledge model with uIV*). Our results showed that the effect of adding a random IV depends on the level of explained variance, *i.e*. in our case on the number of observed dimensions.

When structured IV accounted for less than 50% of the total individual variation, the addition of a random unstructured variation increased community diversity in our simulations. This positive effect was due to the inversions in competitive hierarchy produced by adding a random variation of relatively high variance to individual performance; it allowed more species to be maintained in the community although there were few theoretical winners (*i.e*. species that are the best performing somewhere in the landscape) because of the projection of their niches on few environmental dimensions. Similarly, when the proportion of structured IV was low, adding unstructured IV increased the similarity of the simulated final species abundances with the one of the reference communities. This increase in similarity was however for a great part due to the higher number of species reached when adding unstructured IV (the higher number of zero abundances with the *Imperfect knowledge models without uIV* decreases the estimated similarity with the abundances obtained with the *Perfect knowledge model*).

When the proportion of structured IV increased, this positive effect of adding random IV on community diversity vanished and was even reversed (but see in case of alternative mortality implementation, Appendix 1). This is because the destabilisation of the niche partitioning between species due to unstructured IV - decreased. Indeed, as expected, the lower unstructured IV was (*i.e*. the higher the number of observed dimensions), the greater community mean performance (*i.e*. site sorting) was in comparison to the communities simulated without unstructured IV (see Appendix 3 for further explanation on the absolute differences in community mean performance). This negative effect first increased but then decreased with the number of observed environmental dimensions, because the magnitude (and therefore the effect) of the added unstructured IV became lower. Finally, adding unstructured IV in models is most likely to move simulated community composition away from the reference (here represented by the so-called *Perfect knowledge model*), because this type of variation blurs the species differences that are (although imperfectly) captured with the observed dimensions. In other words, adding randomness does not compensates for lack of knowledge and can even blur the limited knowledge obtained from field data, although this is not the case at a very low level of knowledge of the environment.

Previous modelling studies that tested the effect of adding intraspecific variability on species coexistence provided contrasting results (Lichstein et al. 2007; Vieilledent et al. 2010; Courbaud et al. 2012; Hart et al. 2016; Uriarte and Menge 2018; Crawford et al. 2019). Stump et al. (2022) proposed a framework to explain part of these discrepancies, by differentiating the nature of the traits - niche *vs* hierarchical traits – on which variation was added. While our virtual experiment only considered additional variability in a hierarchical trait (performance) *sensu* Stump et al. (2022), our results here evidenced an additional source of discrepancies when testing the effect of adding a random variability on community dynamics: the relative importance of explained and structured *vs* unexplained and unstructured individual variance. Overall both features, the nature of the traits and its link with performance on the one hand, and its structure or source of variation on the other hand, can explain these contrasting results. Future studies should thus pay great attention to each of these aspects when testing its effect on communities and move away from the systematic approach of adding an unstructured noise.

### Accounting for a high-dimensional environment in community dynamics models

Most previous modelling studies have modelled IV as a random noise around species means (Lichstein et al. 2007; Vieilledent et al. 2010; Courbaud et al. 2012; Hart et al. 2016; Uriarte and Menge 2018; Crawford et al. 2019), and did not represent environmental variations that generate individual variation (*e.g*. Lichstein et al. 2007; Courbaud et al. 2012), or did so in a way that does not mirror multidimensional variation: Uriarte and Menge (2018) provided two different habitats, Vieilledent et al. (2010) used site effects at a much larger scale than individuals, Crawford et al. (2019) represented biotic interactions with resources that are constant through space and time, and while Banitz (2019) is the first to test the consequences of IV resulting from a spatially-structured environmental index, coexistence relied on trade-offs and random disturbances in a one-dimensional environment. Our results, although inevitably dependent on some modelling choices, provided evidence that using independent random draws is not a suitable approach to represent environmentally-driven intraspecific variability in most cases (Girard-Tercieux et al. 2023). To do so, environment-species interactions should be better taken into account in models.

### Improving the knowledge of the environment: a costly but worthy endeavour

The environment can vary in many ways, even if the number of resources is limited, as it is likely the case (Craine 2009). Indeed, many other biotic and abiotic variables can influence the ability to use available resources and individual performance *(e.g*. soil microbiome and texture, microclimate, pathogens, Fortunel et al. 2018; Averill et al. 2022). Moreover, species can partition the same environmental variable (*e.g*. light) by responding non-linearly to it (*e.g*. with different light-performance slopes at different light levels), further increasing the dimensionality of their responses to environmental variation in space and time. As monitoring environmental variables and species responses at fine spatio-temporal scales remains difficult and costly despite technological advances and continuous effort in the field (Estes et al. 2018), part of the environmental variation that influences individuals’ attributes is typically not properly measured in ecological studies.

Our results suggest that improving the characterisation of environmental variation by monitoring additional independent environmental variables (*i.e*. moving to the right in Fig. 2, 3, and 4) is a worthy endeavour. Using one dimension out of 15, 41% of the variation in individual performance is accounted for. The corresponding simulated communities, in absence of any additional random variation, reached less than half the species richness of the communities simulated with the actual 15-dimensional individual performance (median of 4 *vs* 18, Fig. S4.1) with relatively dissimilar community composition (median of similarity in abundance of 0.43 *vs* 0.95, median of mean performance of 1.15 *vs* 1.54, Fig. S4.2). Adding a second dimension allowed to increase the proportion of explained variance in individual performance to 46%, and simulated species richness to a median of 7, with more realistic communities (median of similarity in abundances of 0.59 and median of mean performance of 1.19). The identification of the most influential environmental variables or dimensions in species responses using ecological knowledge (Rüger et al. 2009; Bartlett et al. 2016; Soong et al. 2020) is of course valuable to optimise and prioritise these efforts in the field.

Another way to improve the characterisation of the environment could be to better capture the spatio-temporal structure of the already monitored variables (Tymen et al. 2017; Estes et al. 2018; Zellweger et al. 2019; De Frenne et al. 2021), *i.e*. to monitor them at finer scales in space and time. In our simulation, where we focused on spatial variation, the scale of the environmental variation was the same as the individual (prescribed by the grid mesh size) across all models of individual performance. Testing the effect of degrading the resolution of the observed environmental variation in the case of an imperfect characterisation of the environment could be explored in the future. Finally, improving the characterisation of species responses to a few major environmental variables can also enable to better reveal the realised niche partitioning operating within communities. While niche partitioning is more easily achieved with a high number of environmental dimensions, high level of coexistence can also be reached with only one axis if it is well partitioned among species (*e.g*. Hurtt and Pacala 1995; Detto et al. 2022), thus building a high-dimensional space where each species can perform better somewhere. This is in agreement with several studies that showed significant improvement in the similarity of simulated communities with the reference by only adding a second dimension to species responses in community models (Falster et al. 2017; Rüger et al. 2020).

### Structuring variation: a first step towards accurate representation of multidimensionality

Our virtual experiment builds on an extreme case in which conspecific individuals have exactly the same response to environmental variation and where performance is completely determined by environmental factors, which is unlikely to be the case under the joint effect of environmental and genetic variation in the field, as well as the effect of neutral mechanisms. Partitioning observed IV between genetically-driven, environmentally-driven and unexplained IV using existing data, would be a first step to better understand the nature of IV and to provide hypotheses regarding the resulting structure of IV. This is the goal of many G × E studies and meta-analyses encompassing several ecosystems *(e.g*. Nicotra et al. 2010; Napier et al. 2023). However, while the intraspecific variation that is added in models as a noise around species means is not structured in space and time, IV, whether it is environmentally- or genetically-driven or both, is actually highly likely to be structured in space and time (Girard-Tercieux et al. 2023). This structure could appear in space when IV results from spatially-structured environmental variables or from limited dispersion or local adaptation (Marrot et al. 2021; Schmitt et al. 2021; Westerband et al. 2021). As shown here for spatial variation, this has profound consequences on the properties of the simulated community. Importantly, whatever its source, the spatio-temporal structure of individual variation is an emergent property of conspecific individuals responding more similarly to the environment than heterospecifics locally (Clark 2010; Girard-Tercieux et al. 2023), an important condition for stable species coexistence (Chesson 2000).

Observed or inferred IV, whatever its source (genetic, environmental or an interaction of both, Westerband et al. 2021), can be structured at the individual scale (”individual variability”) using individual effects when one individual is repeatedly observed at one site (Clark et al. 2003). Such individual effects are then typically randomly attributed to individuals in the landscape however (*e.g*. Clark et al. 2007), which is almost equivalent to adding a random noise. Alternatively, the spatial structure of individual effects could be conserved when injected in a model of community dynamics so that a part of observed IV is spatially structured. Pioneer studies have started to explore some aspects of the spatial structure of IV (Purves and Vanderwel 2014; Uriarte and Menge 2018; Banitz 2019), and future work should further explore this direction to generalise its use in community dynamics models. Another source of environmental variation that was not tackled in this study is temporal variation. This variation is often structured, at different temporal scales (seasons, years, El Nin*ñ*o/La Nin*ñ*a events, *etc*.) and this structure should be accounted for in models by expliciting those temporal scales after detection in the data.

Overall, our results suggest that it is crucial to explore the structure of observed IV in real communities to better understand its impact on diversity and community dynamics.

## Acknowledgments

This paper is a joint effort of the working group “INTRACO” supported by CESAB, the synthesis center of the French Foundation for Research on Biodiversity (FRB) and sDiv, the synthesis center of iDiv (DFG FZT 118, 202548816). CGT’s work is supported by a PhD grant provided by the Laboratoire d’Excellence CEBA (Center for the study of Biodiversity in Amazonia; http://www.labex-ceba.fr/en/). CEBA is funded by an “Investissement d’Avenir” grant of the French National Research Agency (CEBA: ANR-10-LABX-25-01). This work has been realised with the support of MESO@LR-Platform at the University of Montpellier, and authors thank particularly Thomas Arsouze and Philippe Verley for their support in the use of the platform.

## Supplementary information and code access

<u>Appendix</u> 1: Alternative implementations of mortality and fecundity

<u>Appendix</u> 2: Stability of the simulations

<u>Appendix</u> 3: Role of suboptimal species depending on the implementation of mortality and fecundity

<u>Appendix</u> 4: Comparisons between communities simulated with the *Imperfect knowledge models without uIV* and with the *Perfect knowledge model*

<u>Appendix</u> 5: Spatial illustration of the experiment

The code used for this study is available in a GitHub repository (https://github.com/camillegirardtercieux/coexist) and has been permanently archived on Zenodo (https://doi.org/10.5281/zenodo.6929042).

## Statements of author roles

CGT, IM and GV conceived the initial ideas and coordinated the INTRACO working group. All authors contributed to the study design and ideas within the INTRACO working group. CGT led the analyses. CGT, IM and GV wrote the first draft of the manuscript, and all authors contributed substantially to revisions.

