## Appendix 1-5 for "Beyond variance: simple random distributions are not a good proxy for intraspecific variability in systems with environmental structure"

### 1 Alternative implementations of mortality and fecundity

To test the robustness of our results to the choices made in building the community dynamics simulator, we implemented alternative ways to simulate mortality and fecundity. For mortality, we explored the three following approaches: (i) the one percent less performing individuals in the landscape die at each timestep, henceforth denoted *deterministic mortality*; (ii) one percent of the individuals die at each timestep, and the probability of each individual to die is proportional to its performance, henceforth denoted *stochastic mortality*; (iii) the probability $\theta_{ij}$ of each individual $j$ to die is computed as a function of its performance, $\theta_{ij} = \text{logit}^{-1}(\text{logit}(0.01) - 0.5 \times p_{ij})$, henceforth denoted *logistic stochastic mortality*. Death events are then drawn in a binomial distribution $B(n_s, \theta)$ with $\theta$ the vector of all $\theta_{ij}$. For fecundity, we explored the two following approaches: (i) the number of propagules $prop_{j,t}$ depends on species abundance $A_{j,t}$: $prop_{j,t} = round(0.5 \times A_{j,t})$, henceforth denoted the *abundance-dependent fecundity*; or (ii) each species present in the community produces ten offspring per timestep, henceforth denoted the *fixed fecundity*. Results obtained with *deterministic mortality* and *abundance-dependent fecundity* are presented in the main text, and we present below the results with other mortality and fecundity alternatives (Fig. S1.1 to Fig. S1.10).

#### *Deterministic mortality* and *fixed fecundity*

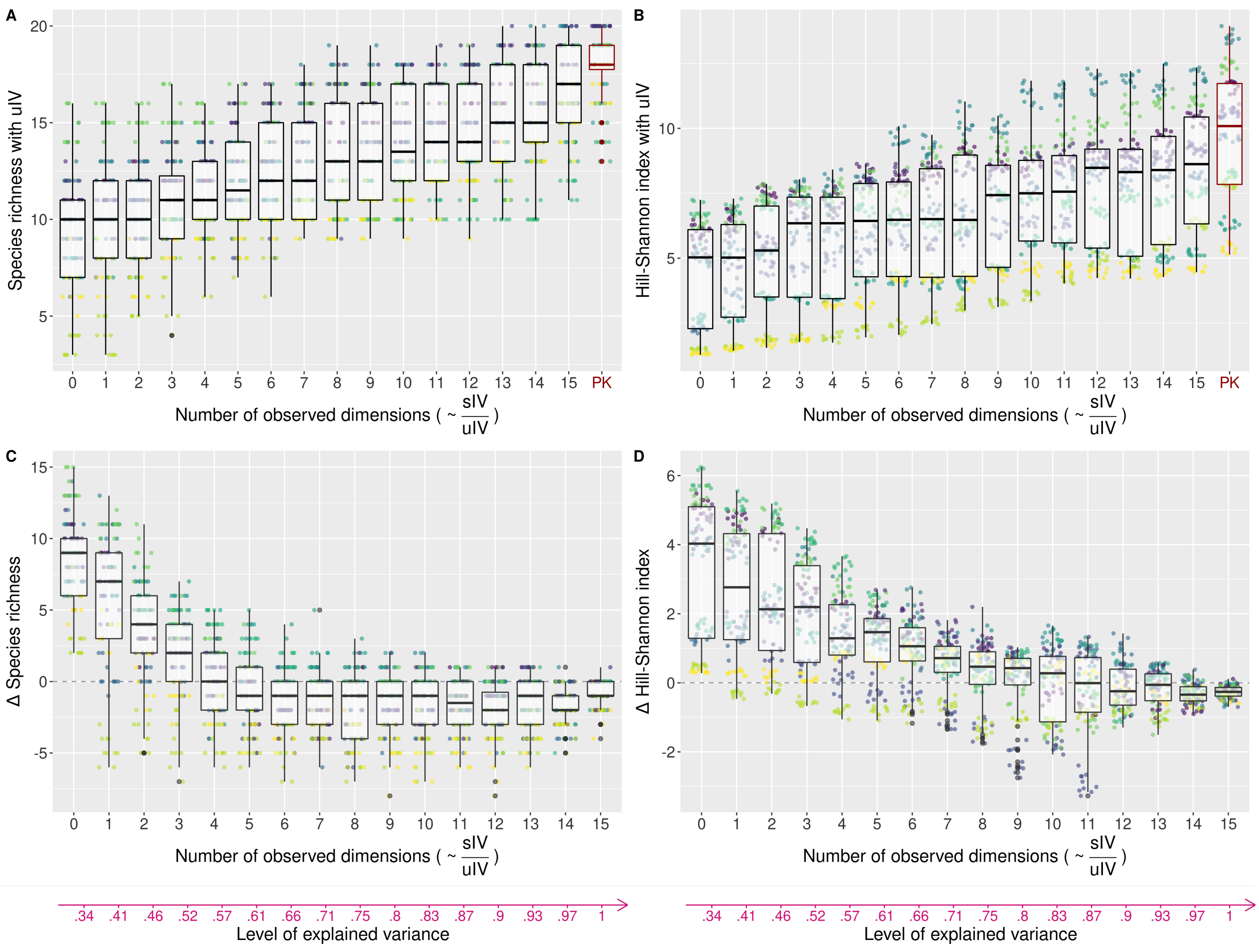


**Figure S1.1: Effect of the structure of individual variation on community diversity.** Each point represents the diversity, either computed as the species richness – left panels – or the Hill-Shannon diversity index – right panels – of a final simulated community. Each color represents an $E \times S$ configuration (ten points per color, for the ten initial conditions). The horizontal axis corresponds to the number of observed environmental dimensions, which is proportional to the ratio of structured and unstructured IV in the performance models. Each number of observed dimensions corresponds to a level of explained variance in individual performance (see Fig. 2 of main text) depicted with the pink arrow at the bottom. The top panels show the final community diversity obtained with the *Imperfect knowledge models with uIV* (0 to 15 observed dimensions) and with the *Perfect knowledge model* (PK, red, far right). The bottom panels show the difference in the final community diversity obtained with the *Imperfect knowledge models with* and *without uIV*. Points that are above zero (horizontal dashed line) correspond to a higher diversity when adding unstructured IV. Results shown here were obtained with a *deterministic mortality* and a *fixed fecundity*.

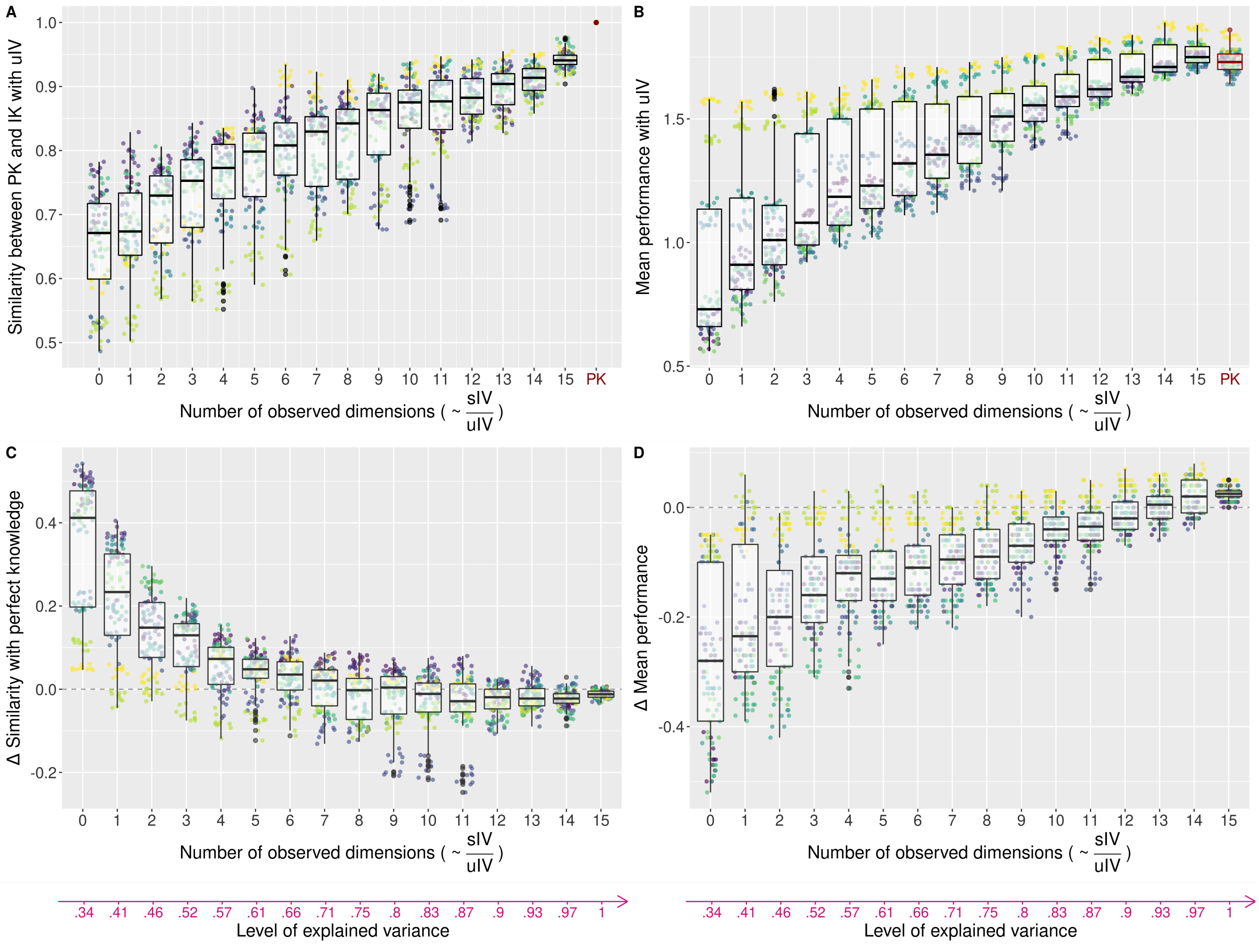


**Figure S1.2: Effect of the structure of individual variation on the similarity in final species abundances between models and on the of the final communities.** Each color represents an $E \times S$ configuration. For the similarity - left panels -, each point represents the pairwise percentage similarity (PS) in the final species abundances between two simulations with the same $E \times S$ configuration and the same initial conditions (ten points per color), but obtained using the *Perfect knowledge model* one the one hand and one of the *Imperfect knowledge models* on the other hand. For the site sorting - right panels -, each point represents the community mean performance of the final communities. This mean performance was calculated with the *Perfect knowledge model* and averaged across all individuals at the end of the simulation. The top panels show these two metrics for communities simulated with the *Imperfect knowledge models with uIV* (0 to 15 observed dimensions) and with the *Perfect knowledge model* (PK, red, far right). The bottom panels show the difference in these metrics for communities obtained with the *Imperfect knowledge models with* and *without uIV*. Points that are above zero (horizontal dashed line) correspond to a higher similarity or mean performance when adding unstructured IV, respectively. Results shown here were obtained with a *deterministic mortality* and a *fixed fecundity*.

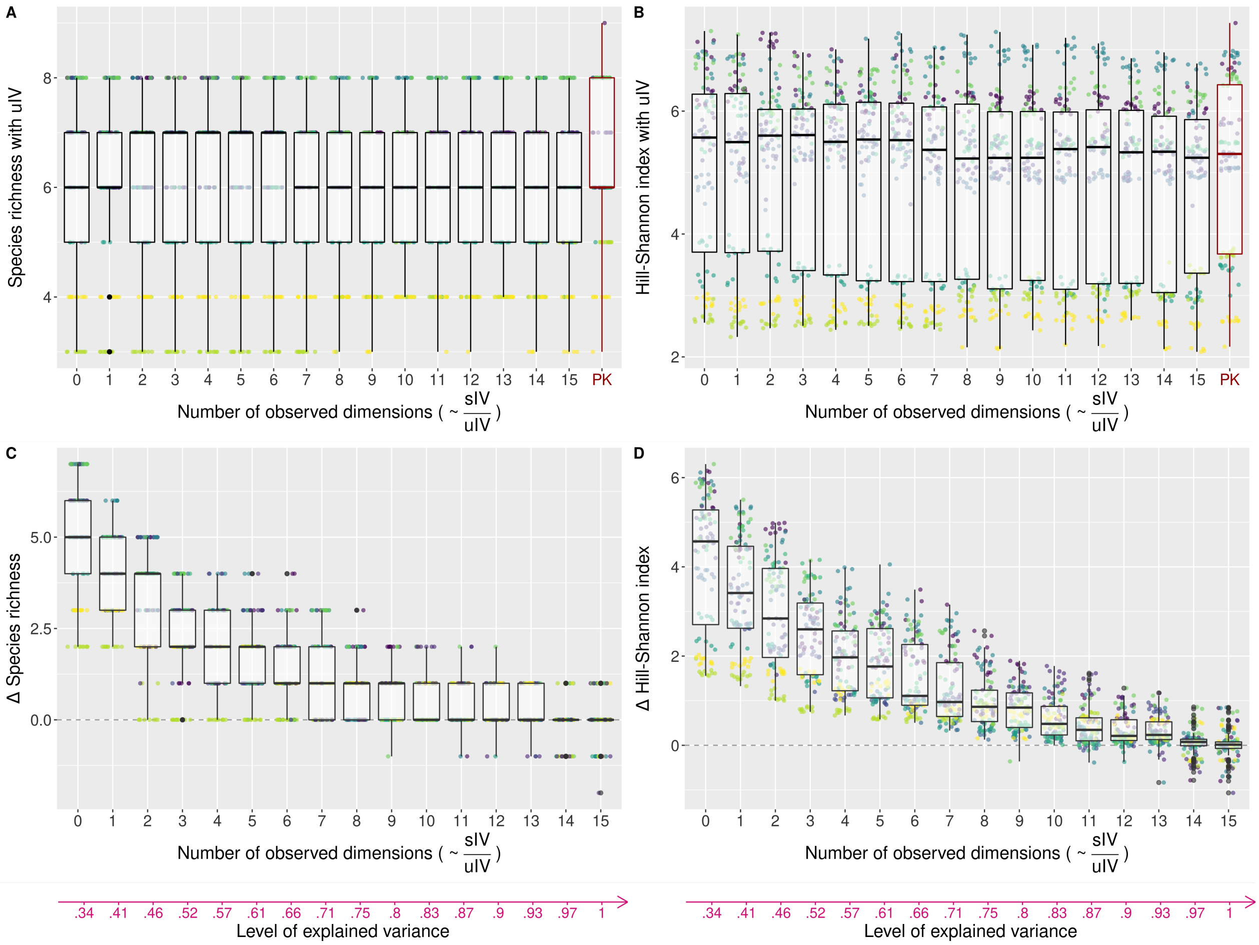


**Figure S1.3: Effect of the structure of individual variation on community diversity.** Each point represents the diversity, either computed as the species richness – left panels – or the Hill-Shannon diversity index – right panels – of a final simulated community. Each color represents an $E \times S$ configuration (ten points per color, for the ten initial conditions). The horizontal axis corresponds to the number of observed environmental dimensions, which is proportional to the ratio of structured and unstructured IV in the performance models. Each number of observed dimensions corresponds to a level of explained variance in individual performance (see Fig. 2 of main text) depicted with the pink arrow at the bottom. The top panels show the final community diversity obtained with the *Imperfect knowledge models with uIV* (0 to 15 observed dimensions) and with the *Perfect knowledge model* (PK, red, far right). The bottom panels show the difference in the final community diversity obtained with the *Imperfect knowledge models with* and *without uIV*. Points that are above zero (horizontal dashed line) correspond to a higher diversity when adding unstructured IV. Results shown here were obtained with a *stochastic mortality* and an *abundance-dependent fecundity*.

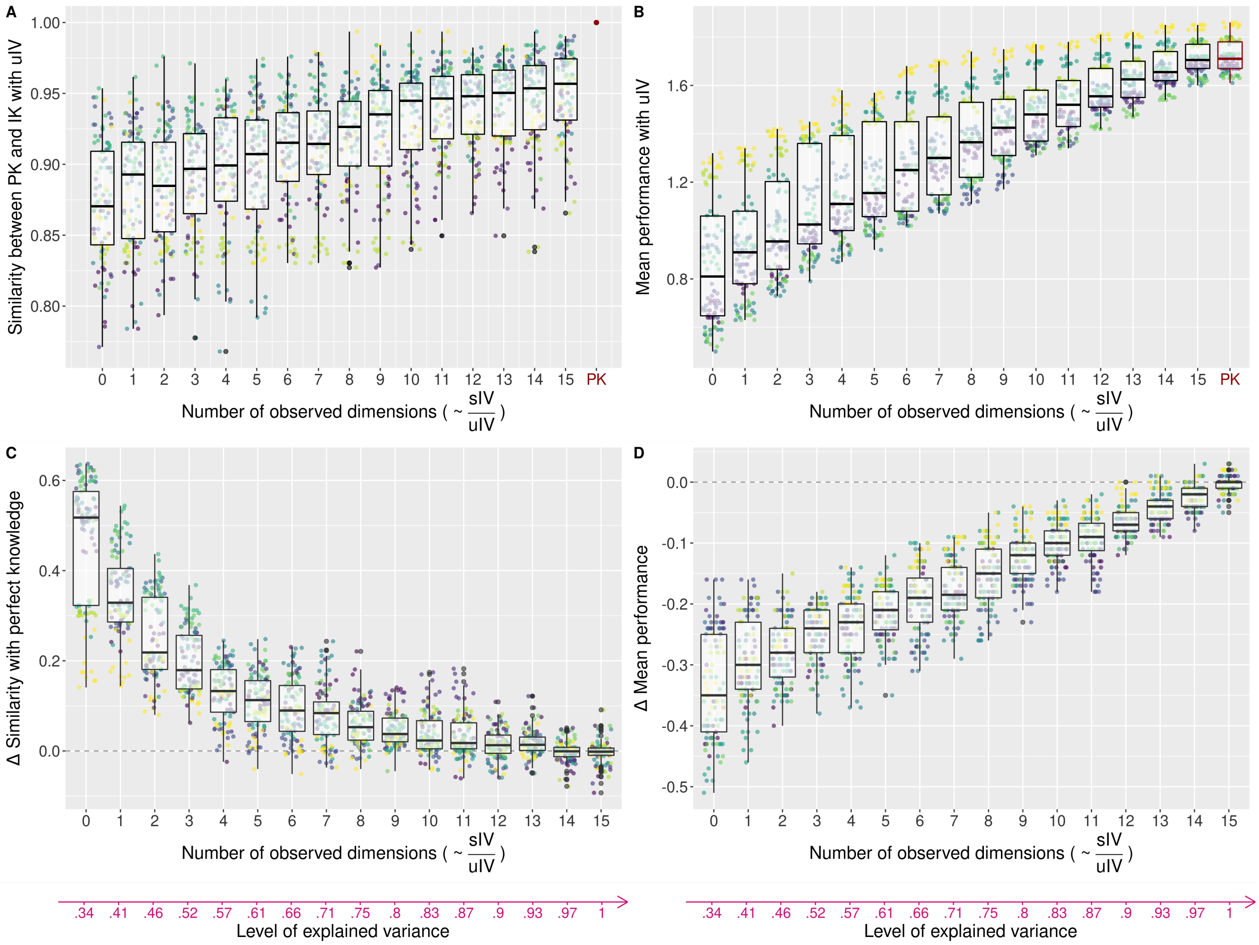


**Figure S1.4: Effect of the structure of individual variation on the similarity in final species abundances between models and on the of the final communities.** Each color represents an $E \times S$ configuration. For the similarity - left panels -, each point represents the pairwise percentage similarity (PS) in the final species abundances between two simulations with the same $E \times S$ configuration and the same initial conditions (ten points per color), but obtained using the *Perfect knowledge model* one the one hand and one of the *Imperfect knowledge models* on the other hand. For the site sorting - right panels -, each point represents the community mean performance of the final communities. This mean performance was calculated with the *Perfect knowledge model* and averaged across all individuals at the end of the simulation. The top panels show these two metrics for communities simulated with the *Imperfect knowledge models with uIV* (0 to 15 observed dimensions) and with the *Perfect knowledge model* (PK, red, far right). The bottom panels show the difference in these metrics for communities obtained with the *Imperfect knowledge models with* and *without uIV*. Points that are above zero (horizontal dashed line) correspond to a higher similarity or mean performance when adding unstructured IV, respectively. Results shown here were obtained with a *stochastic mortality* and an *abundance-dependent fecundity*.

#### ***Stochastic mortality*** **and *fixed fecundity***

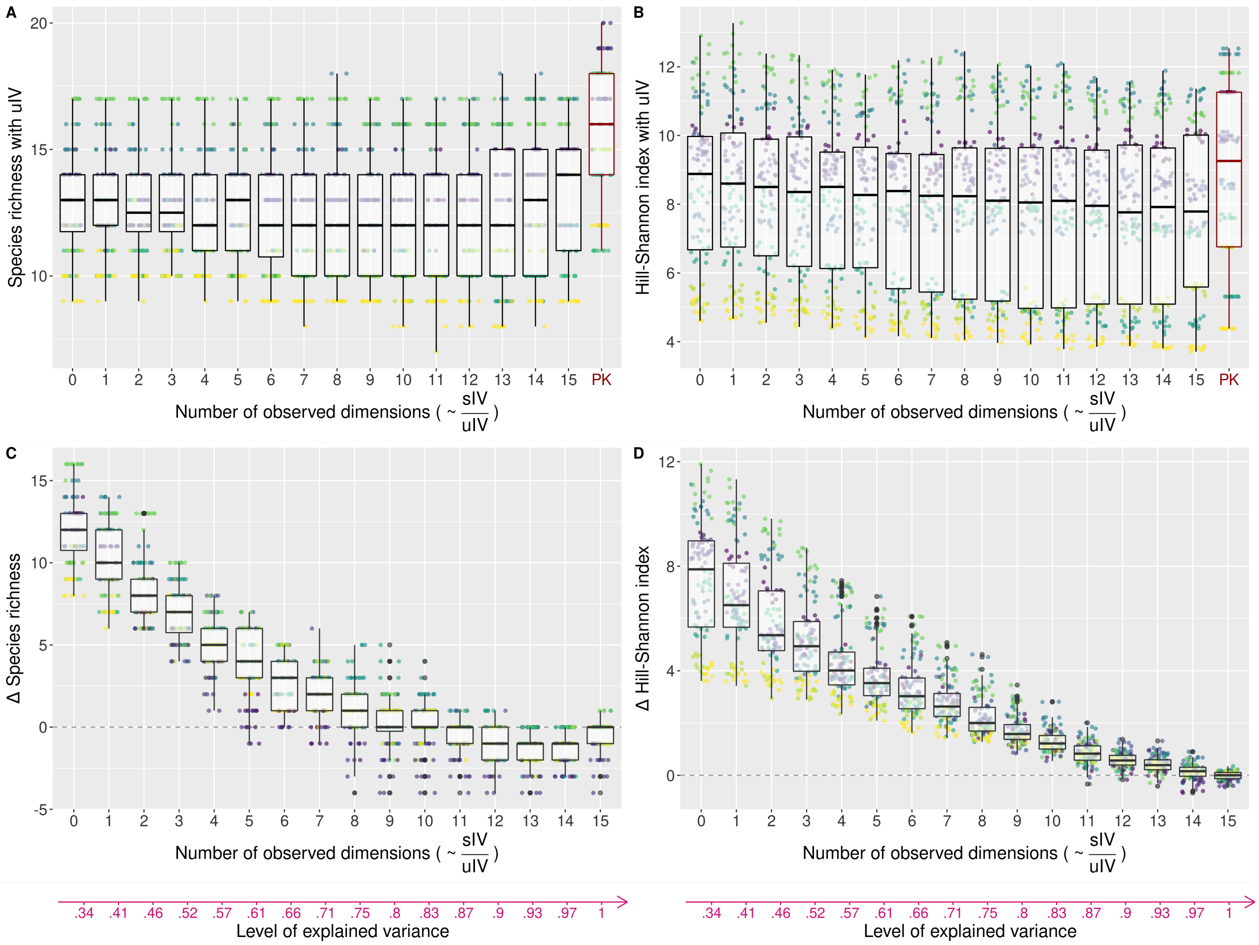


**Figure S1.5: Effect of the structure of individual variation on community diversity.** Each point represents the diversity, either computed as the species richness – left panels – or the Hill-Shannon diversity index – right panels – of a final simulated community. Each color represents an $E \times S$ configuration (ten points per color, for the ten initial conditions). The horizontal axis corresponds to the number of observed environmental dimensions, which is proportional to the ratio of structured and unstructured IV in the performance models. Each number of observed dimensions corresponds to a level of explained variance in individual performance (see Fig. 2 of main text) depicted with the pink arrow at the bottom. The top panels show the final community diversity obtained with the *Imperfect knowledge models with uIV* (0 to 15 observed dimensions) and with the *Perfect knowledge model* (PK, red, far right). The bottom panels show the difference in the final community diversity obtained with the *Imperfect knowledge models with* and *without uIV*. Points that are above zero (horizontal dashed line) correspond to a higher diversity when adding unstructured IV. Results shown here were obtained with a *stochastic mortality* and a *fixed fecundity*.

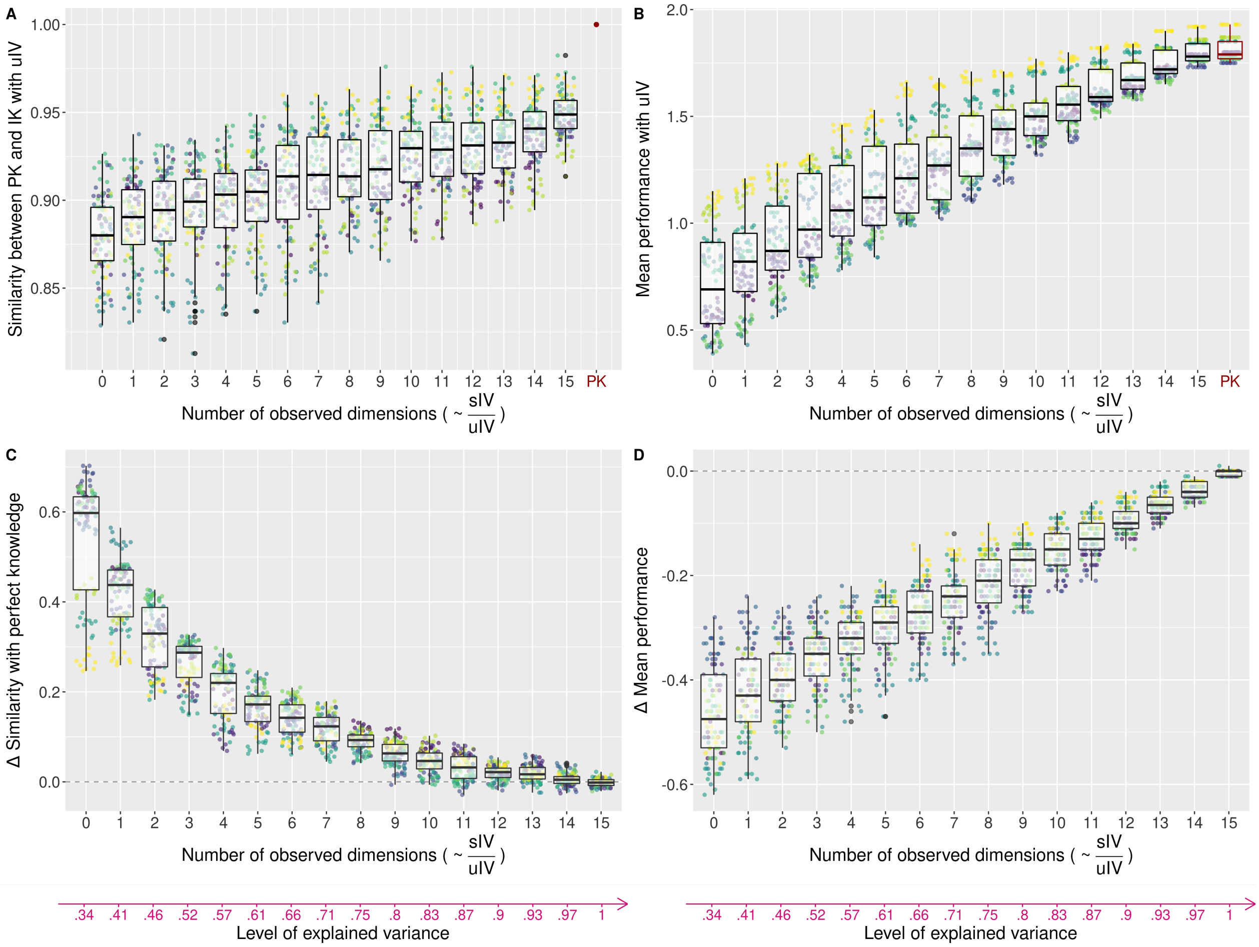


**Figure S1.6: Effect of the structure of individual variation on the similarity in final species abundances between models and on the of the final communities.** Each color represents an $E \times S$ configuration. For the similarity - left panels -, each point represents the pairwise percentage similarity (PS) in the final species abundances between two simulations with the same $E \times S$ configuration and the same initial conditions (ten points per color), but obtained using the *Perfect knowledge model* one the one hand and one of the *Imperfect knowledge models* on the other hand. For the site sorting - right panels -, each point represents the community mean performance of the final communities. This mean performance was calculated with the *Perfect knowledge model* and averaged across all individuals at the end of the simulation. The top panels show these two metrics for communities simulated with the *Imperfect knowledge models with uIV* (0 to 15 observed dimensions) and with the *Perfect knowledge model* (PK, red, far right). The bottom panels show the difference in these metrics for communities obtained with the *Imperfect knowledge models with* and *without uIV*. Points that are above zero (horizontal dashed line) correspond to a higher similarity or mean performance when adding unstructured IV, respectively. Results shown here were obtained with a *stochastic mortality* and a *fixed fecundity*.

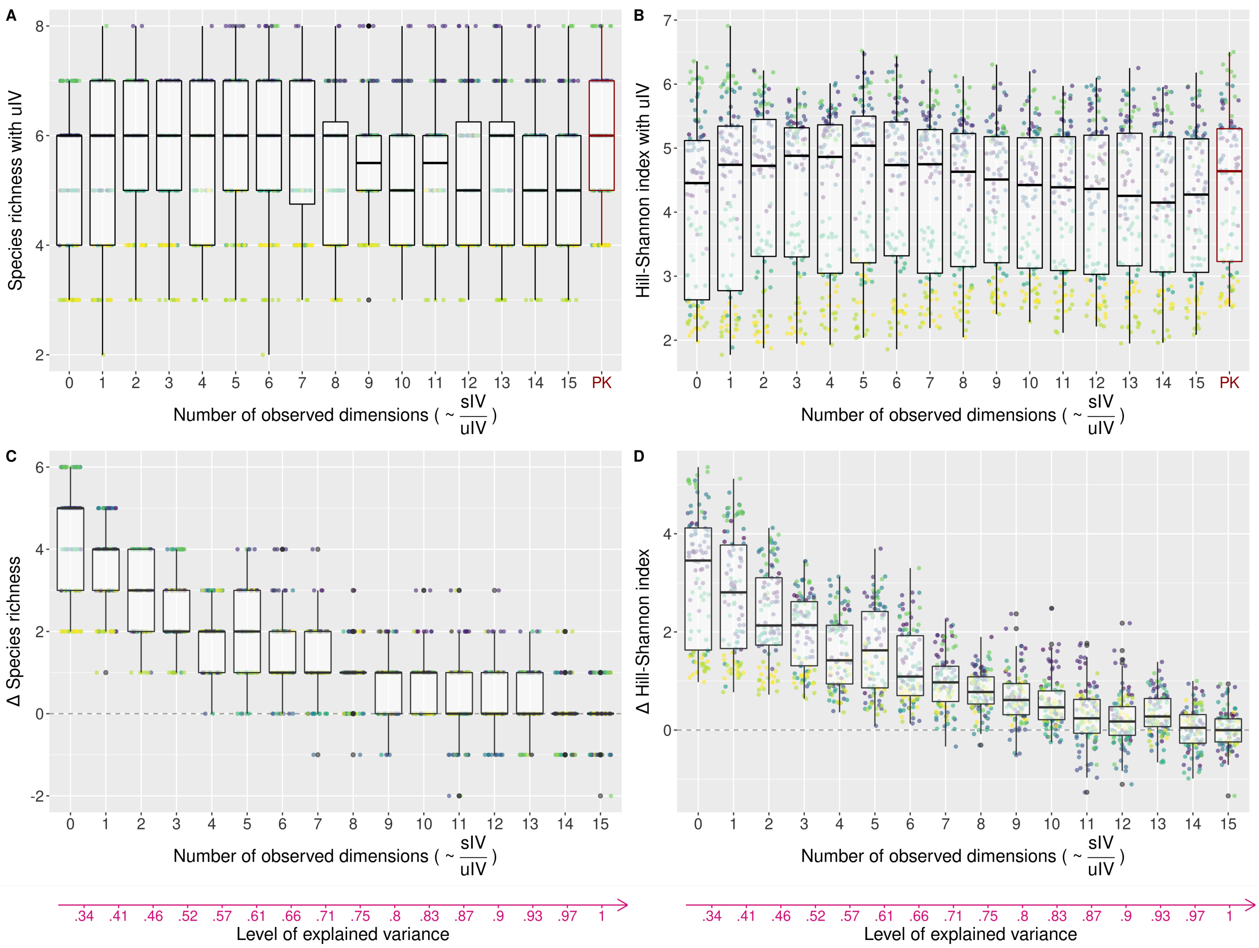


**Figure S1.7: Effect of the structure of individual variation on community diversity.** Each point represents the diversity, either computed as the species richness – left panels – or the Hill-Shannon diversity index – right panels – of a final simulated community. Each color represents an $E \times S$ configuration (ten points per color, for the ten initial conditions). The horizontal axis corresponds to the number of observed environmental dimensions, which is proportional to the ratio of structured and unstructured IV in the performance models. Each number of observed dimensions corresponds to a level of explained variance in individual performance (see Fig. 2 of main text) depicted with the pink arrow at the bottom. The top panels show the final community diversity obtained with the *Imperfect knowledge models with uIV* (0 to 15 observed dimensions) and with the *Perfect knowledge model* (PK, red, far right). The bottom panels show the difference in the final community diversity obtained with the *Imperfect knowledge models with* and *without uIV*. Points that are above zero (horizontal dashed line) correspond to a higher diversity when adding unstructured IV. Results shown here were obtained with a *logistic stochastic mortality* and an *abundance-dependent fecundity*.

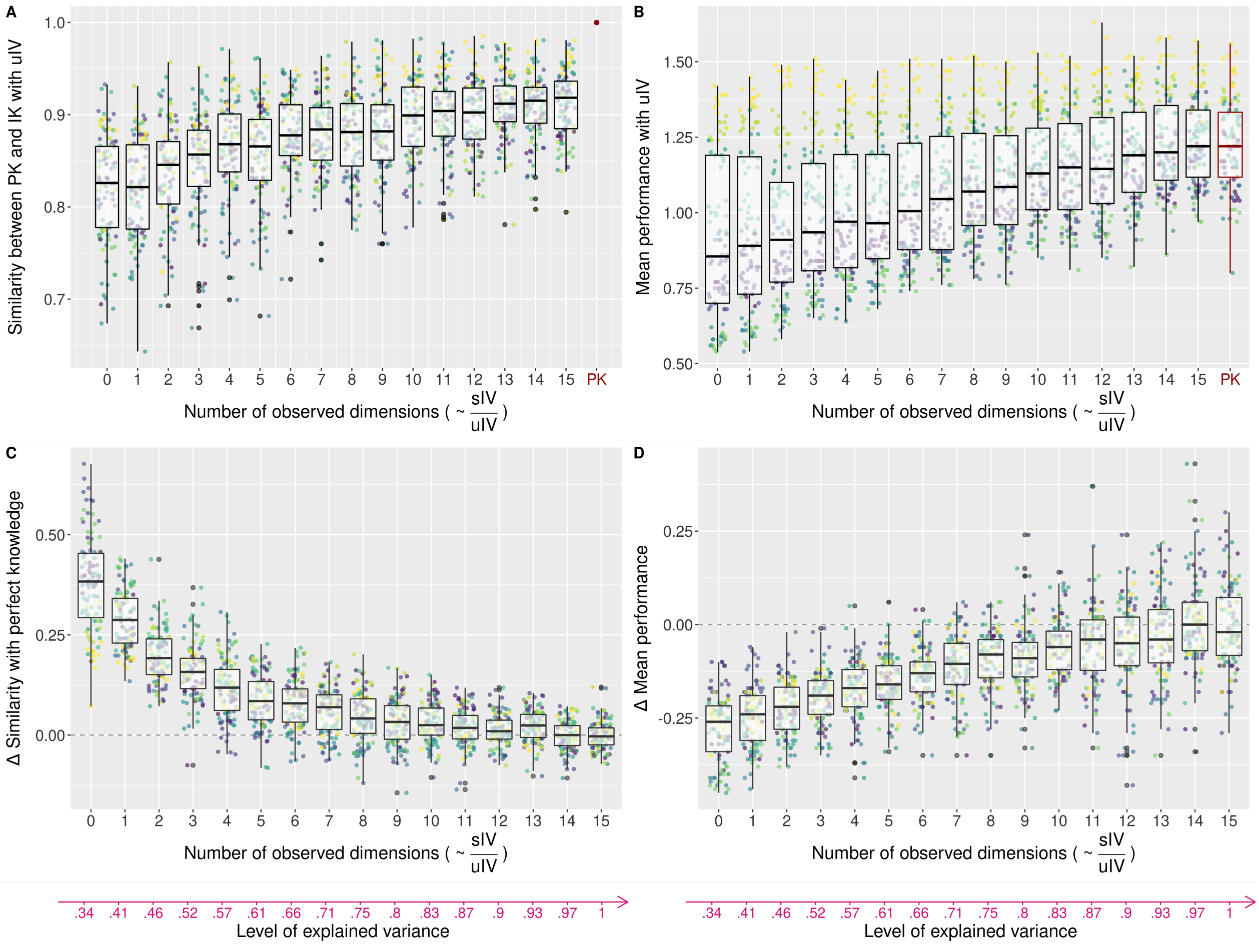


**Figure S1.8: Effect of the structure of individual variation on the similarity in final species abundances between models and on the of the final communities.** Each color represents an $E \times S$ configuration. For the similarity - left panels -, each point represents the pairwise percentage similarity (PS) in the final species abundances between two simulations with the same $E \times S$ configuration and the same initial conditions (ten points per color), but obtained using the *Perfect knowledge model* one the one hand and one of the *Imperfect knowledge models* on the other hand. For the site sorting - right panels -, each point represents the community mean performance of the final communities. This mean performance was calculated with the *Perfect knowledge model* and averaged across all individuals at the end of the simulation. The top panels show these two metrics for communities simulated with the *Imperfect knowledge models with uIV* (0 to 15 observed dimensions) and with the *Perfect knowledge model* (PK, red, far right). The bottom panels show the difference in these metrics for communities obtained with the *Imperfect knowledge models with* and *without uIV*. Points that are above zero (horizontal dashed line) correspond to a higher similarity or mean performance when adding unstructured IV, respectively. Results shown here were obtained with a *logistic stochastic mortality* and an *abundance-dependent fecundity*.

#### *Logistic stochastic mortality* and *fixed fecundity*

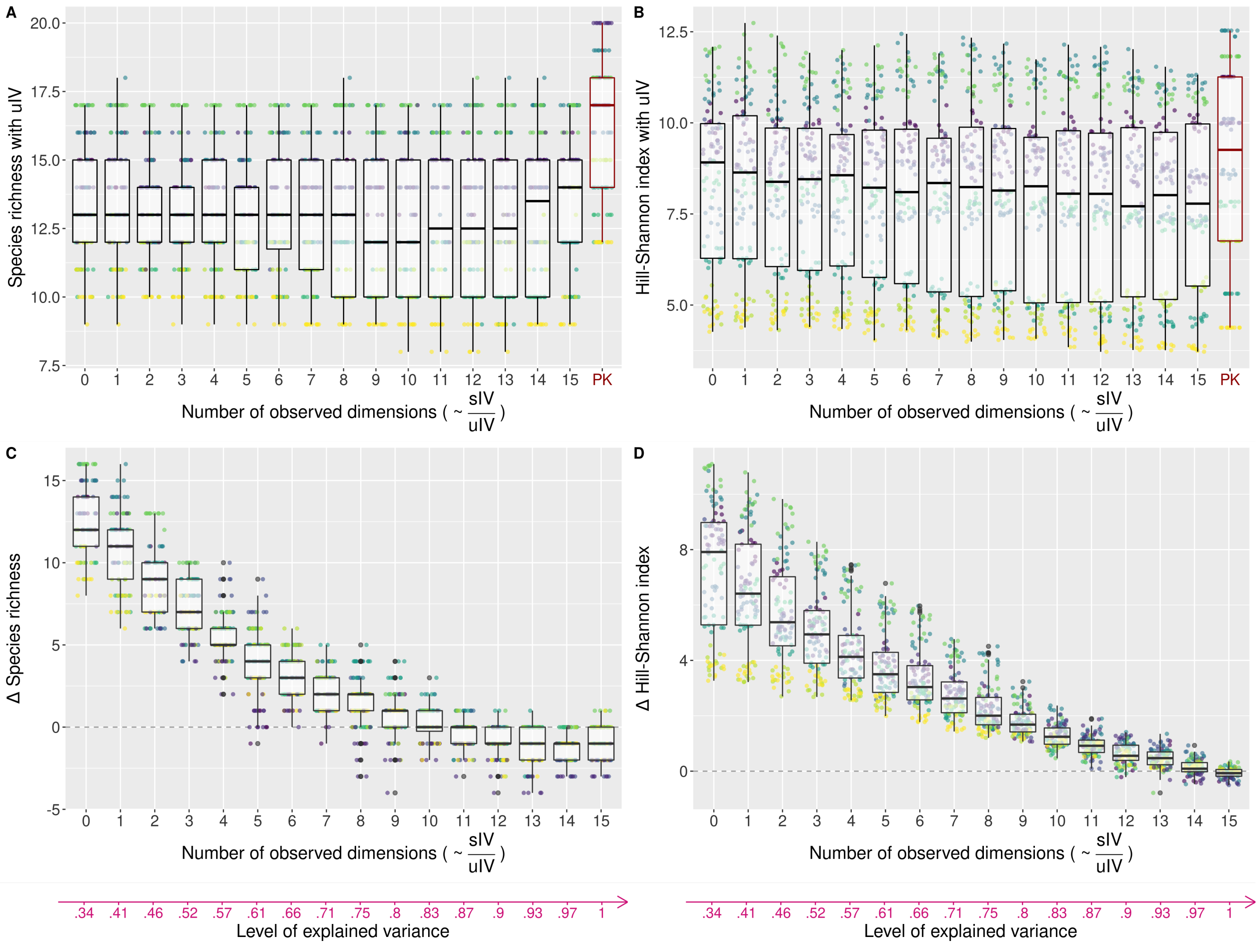


**Figure S1.9: Effect of the structure of individual variation on community diversity.** Each point represents the diversity, either computed as the species richness – left panels – or the Hill-Shannon diversity index – right panels – of a final simulated community. Each color represents an $E \times S$ configuration (ten points per color, for the ten initial conditions). The horizontal axis corresponds to the number of observed environmental dimensions, which is proportional to the ratio of structured and unstructured IV in the performance models. Each number of observed dimensions corresponds to a level of explained variance in individual performance (see Fig. 2 of main text) depicted with the pink arrow at the bottom. The top panels show the final community diversity obtained with the *Imperfect knowledge models with uIV* (0 to 15 observed dimensions) and with the *Perfect knowledge model* (PK, red, far right). The bottom panels show the difference in the final community diversity obtained with the *Imperfect knowledge models with* and *without uIV*. Points that are above zero (horizontal dashed line) correspond to a higher diversity when adding unstructured IV. Results shown here were obtained with a *logistic stochastic mortality* and a *fixed fecundity*.

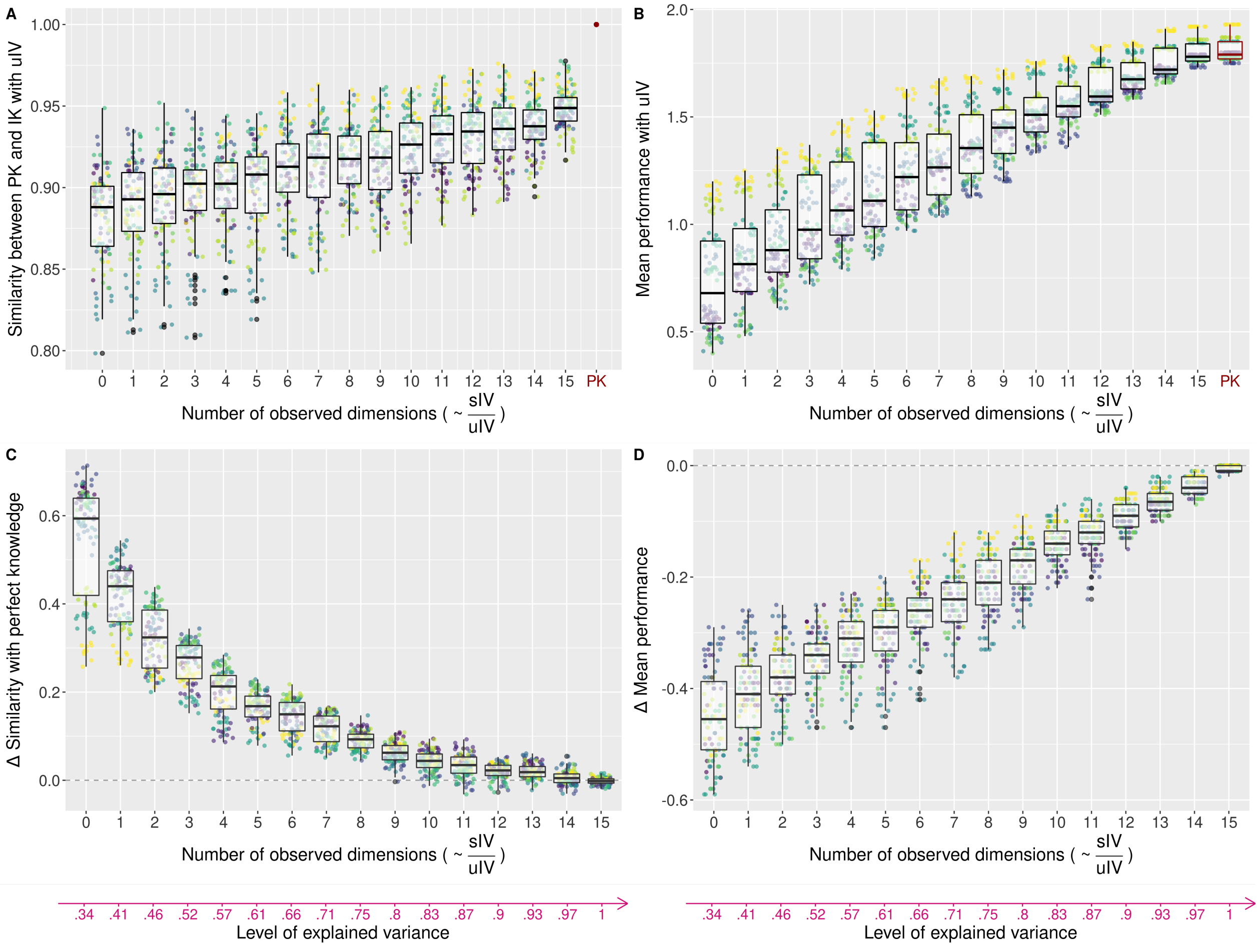


**Figure S1.10: Effect of the structure of individual variation on the similarity in final species abundances between models and on the of the final communities.** Each color represents an $E \times S$ configuration. For the similarity - left panels -, each point represents the pairwise percentage similarity (PS) in the final species abundances between two simulations with the same $E \times S$ configuration and the same initial conditions (ten points per color), but obtained using the *Perfect knowledge model* one the one hand and one of the *Imperfect knowledge models* on the other hand. For the site sorting - right panels -, each point represents the community mean performance of the final communities. This mean performance was calculated with the *Perfect knowledge model* and averaged across all individuals at the end of the simulation. The top panels show these two metrics for communities simulated with the *Imperfect knowledge models with uIV* (0 to 15 observed dimensions) and with the *Perfect knowledge model* (PK, red, far right). The bottom panels show the difference in these metrics for communities obtained with the *Imperfect knowledge models with* and *without uIV*. Points that are above zero (horizontal dashed line) correspond to a higher similarity or mean performance when adding unstructured IV, respectively. Results shown here were obtained with a *logistic stochastic mortality* and a *fixed fecundity*.

### 2 Stability of the simulations

We would expect the simulations made with unstructured IV in individual performance to be characterized by significantly different final communities with various species abundance distributions (unstable coexistence, Hubbell 2001). However, all three models of individual performance produced communities with relatively similar final species abundance distribution, and this stability was actually even lower for the *Perfect knowledge model* (Fig. S2.11).

This was explained by the fact that with the *Perfect knowledge model*, rare species can be maintained in the community but with a high dependence on initial conditions. On the contrary, *Imperfect knowledge models* enable mainly species with abundant suitable habitat to persist in the community, and are therefore less sensitive to initial conditions. With many observed dimensions, the *Imperfect knowledge models without uIV* even become less stable than the *Imperfect knowledge models with uIV*. As the *Imperfect knowledge models without uIV* approach the *Perfect knowledge model*, it enables more rare species to be maintained, with a high dependence on initial conditions.

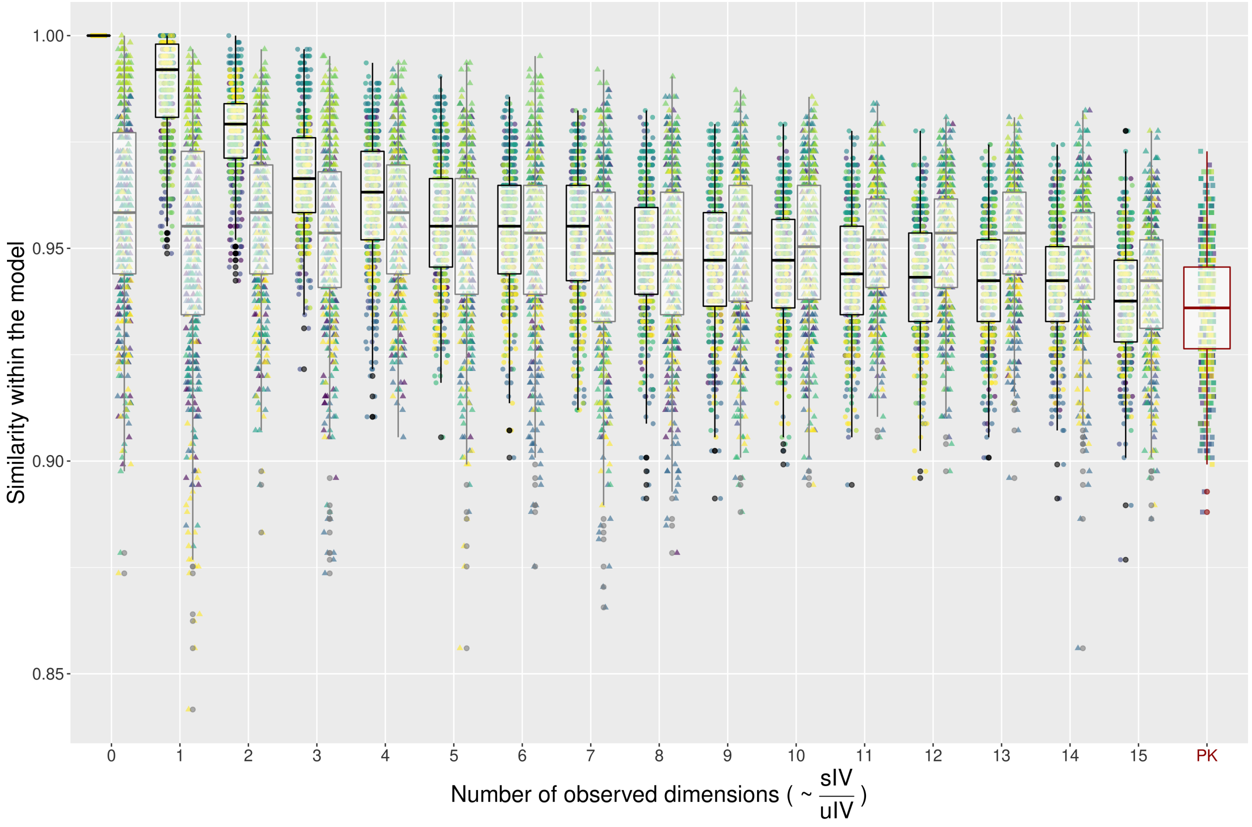


**Figure S2.11: Effect of the structure of individual variation on the stability of the community composition within models.** Each point represents the pairwise percentage similarity of the species abundances of the final community of two repetitions of the same model with the same $E \times S$ configuration. Each color represents a $E \times S$ configuration (45 points per color). The horizontal axis corresponds to the number of observed environmental dimensions, which is proportional to the ratio of structured and unstructured IV in the performance models. At a given number of observed dimensions, the black boxplot on the left corresponds to communities simulated with the *Imperfect knowledge model without uIV* while the gray boxplot on the right corresponds to communities simulated with the *Imperfect knowledge model with uIV*, except on the far right (PK) where the red boxplot, represents communities simulated with the *Perfect knowledge model*. Results shown here were obtained with a *deterministic mortality* and an *abundance-dependent fecundity*.

### 3 Role of suboptimal species depending on the implementation of mortality and fecundity

With some particular environments and species optima ($E \times S$ configuration), some species that are theoretically not winners in the landscape, i.e. that can be outperformed by another species in any site, can be maintained. We refer to these species as suboptimal species. These suboptimal species have optima that are similar to the optimum of a dominant species (or theoretical winner) and their performance at the optimum is therefore high (although suboptimal). Suboptimal species present on sites close to the optimum of an actual dominant species, can have a higher performance than actual dominant species on some other sites.

With *deterministic mortality*, only a determined number of the less performant individuals die at each iteration. Therefore, individuals belonging to dominant species die rather than individuals of suboptimal species, enabling a high level of coexistence even when there are only few theoretical winners. As a result, the communities simulated with the *Perfect knowledge model* (or the *Imperfect knowledge model* with fifteen dimensions) can be outperformed in terms of mean performance because suboptimal species can persist (Fig. 4B of the main text). However, this effect varies depending on the way mortality was implemented.

With *stochastic mortality*, the difference in mean performance between communities simulated with or without unstructured IV tends to zero as unstructured IV decreases. With *deterministic mortality* however, the difference is always higher than with *stochastic mortality*, so that starting from seven observed dimensions, the communities tend to have a higher mean performance with unstructured IV (Fig. 4D of main text). This is due to the number of individuals belonging to suboptimal species. In the case of a *deterministic mortality*, the difference in mean performance between communities simulated with vs. without unstructured IV is tightly negatively related to the difference in the number of individuals belonging to suboptimal species (Fig. S3.12). Indeed, the number of suboptimal individuals is higher with deterministic than *stochastic mortality* since suboptimal individuals can durably persist as never being among the least performing individuals in the community that are filtered out at each time step (Fig. S3.13). Moreover, the addition of unstructured IV impacts more negatively the number of individuals of suboptimal species in the case of deterministic rather than *stochastic mortality* (Fig. S3.14). Indeed, unstructured IV reduces the possibility to generate individuals that durably persists: in average, half of them will have a lower performance with unstructured IV than without, increasing their chance of being filtered out.

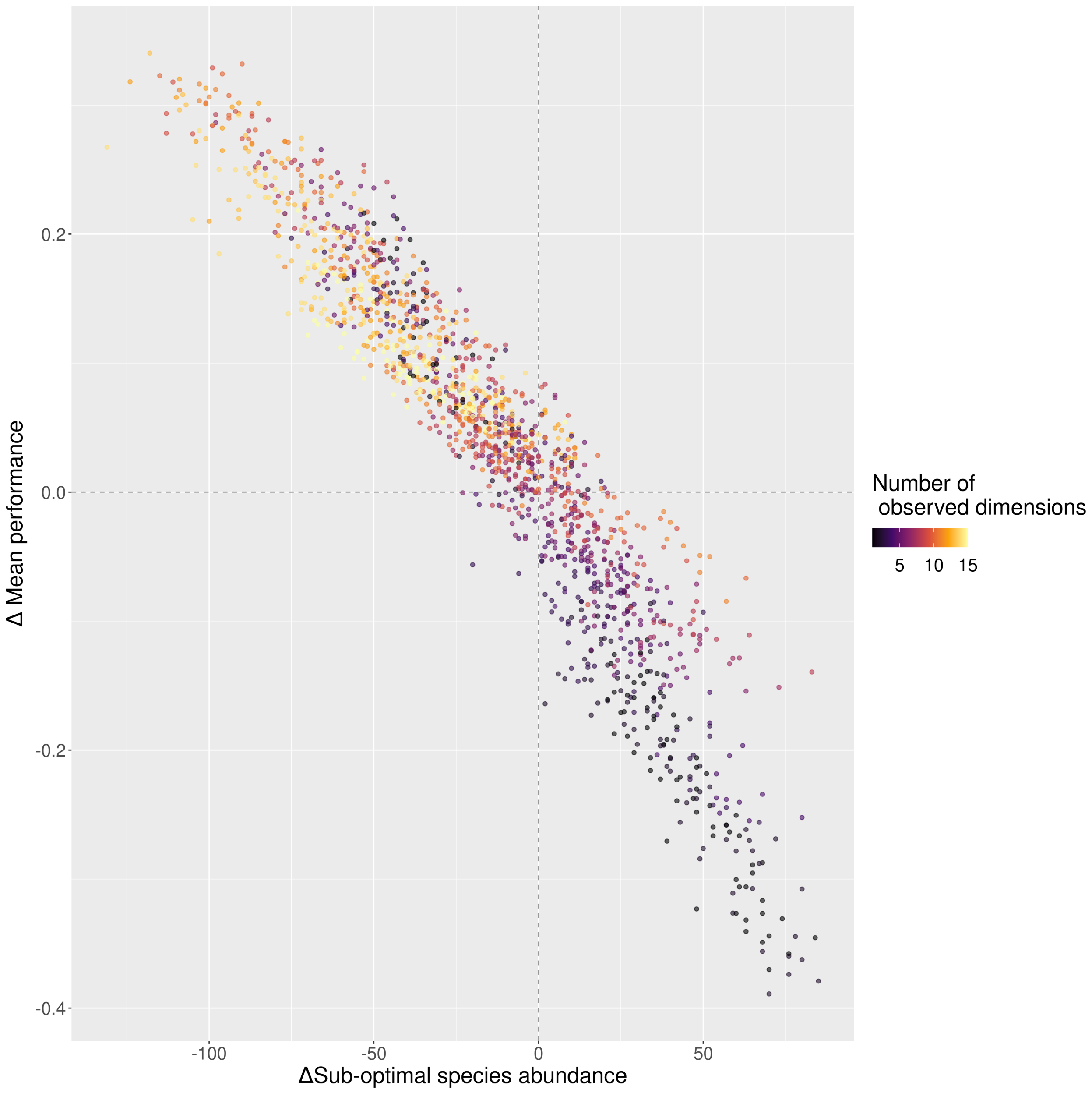


**Figure S3.12: Relationship between the effect of adding unstructured IV on suboptimal species abundances and mean performance.** Results shown here were obtained with a *deterministic mortality* and an *abundance-dependent fecundity*.

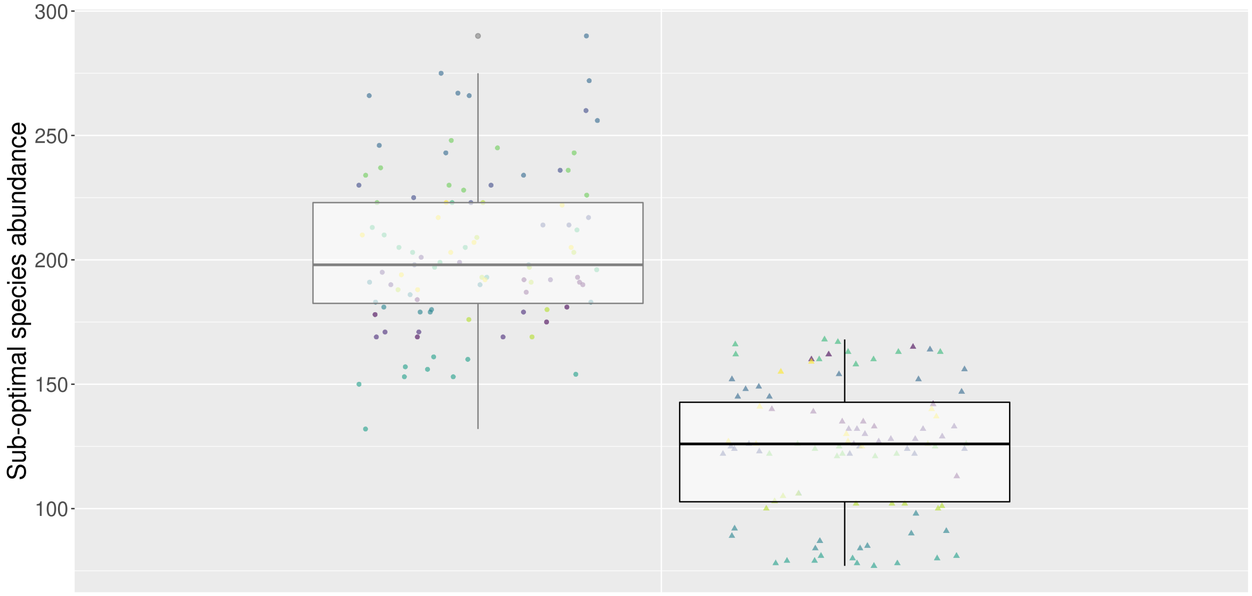


**Figure S3.13: Abundance of suboptimal species in the final communities simulated with deterministic *vs. stochastic mortality.*** Each point represents the total number of individuals of suboptimal species in the final communities obtained with the *Perfect knowledge model*. Each color represents an $E \times S$ configuration (ten points per color, for the ten initial conditions). The gray boxplot on the left corresponds to communities simulated with a *deterministic mortality* while the black boxplot on the right corresponds to communities simulated with a *stochastic mortality*. Results shown here were obtained with *abundance-dependent fecundity*.

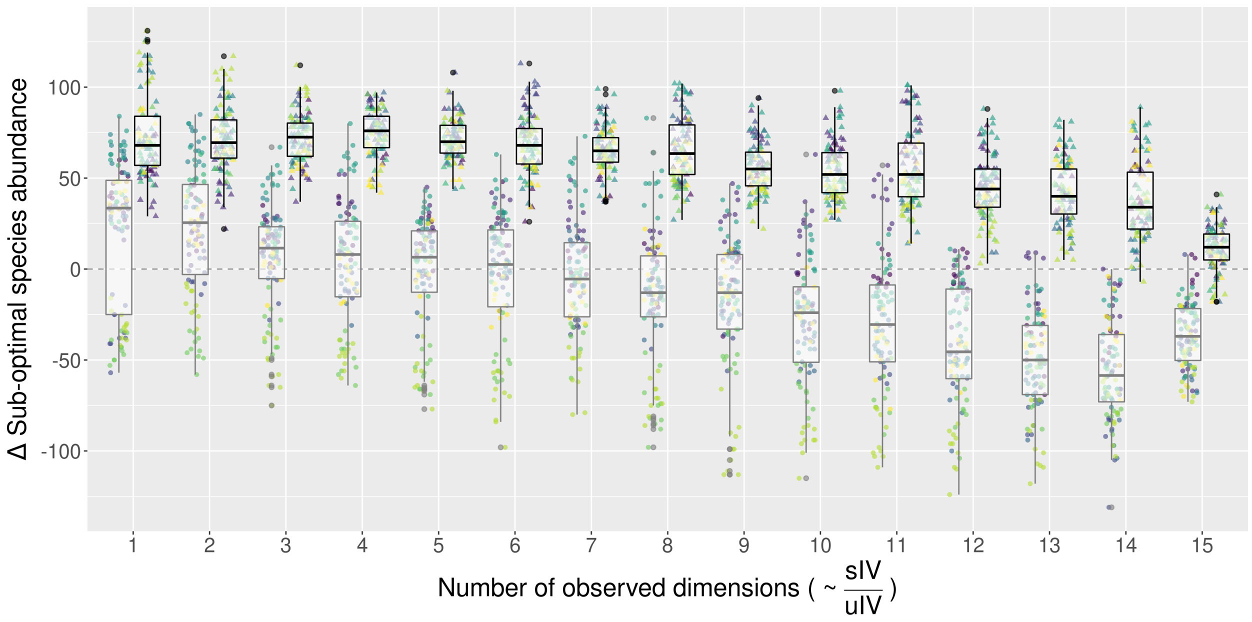


**Figure S3.14: Comparison of the effect of adding unstructured IV on the abundance of suboptimal species between communities simulated with deterministic *vs. stochastic mortality.*** Each color represents an $E \times S$ configuration. Each point represents the difference between the abundance of suboptimal species in the final communities obtained with the same $E \times S$ configuration and the same initial conditions (ten points per color), but simulated with the *Imperfect knowledge models with vs. without uIV*. Points that are above zero (horizontal dashed line) correspond to a higher abundance of suboptimal species when adding unstructured IV. At a given number of observed dimensions, the gray boxplot on the left corresponds to communities simulated with a *deterministic mortality* while the black boxplot on the right corresponds to communities simulated with a *stochastic mortality*. Results shown here were obtained with *abundance-dependent fecundity*.

### 4 Comparisons between communities simulated with the *Imperfect knowledge models without uIV* and with the *Perfect knowledge model*

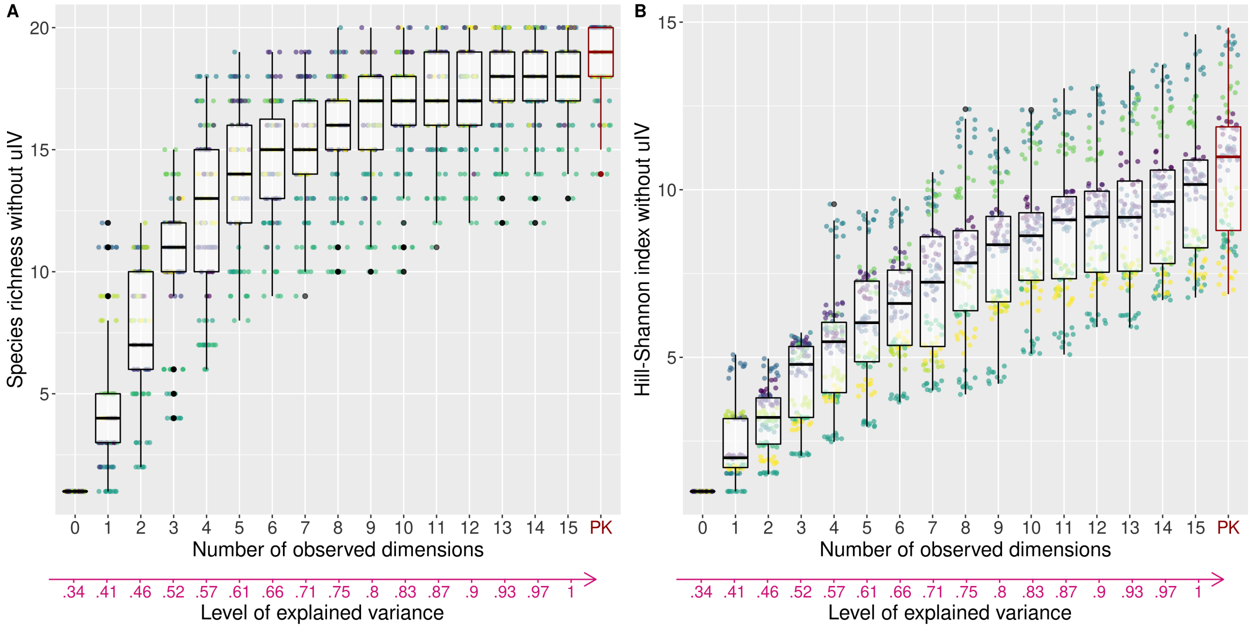


**Figure S4.15: Effect of the number of observed environmental dimensions on community diversity.** Each point represents the diversity, either computed as the species richness (A) or the Hill-Shannon diversity index (B) of a final simulated community. Each color represents an $E \times S$ configuration (ten points per color, for the ten initial conditions). The horizontal axis corresponds to the number of observed environmental dimensions. Each number of observed dimensions corresponds to a level of explained variance in individual performance (see Fig. 2 of main text) depicted with the pink arrow at the bottom. The vertical axis corresponds to the final species richness obtained with the *Imperfect knowledge models without uIV* (0 to 15 observed dimensions) and with the *Perfect knowledge model* (PK, red, far right). Comparing the *Imperfect knowledge models without uIV* with the *Perfect knowledge model* is useful to examine the effect of the reduction of the number of observed dimensions on species richness. Results shown here were obtained with a *deterministic mortality* and an *abundance-dependent fecundity*.

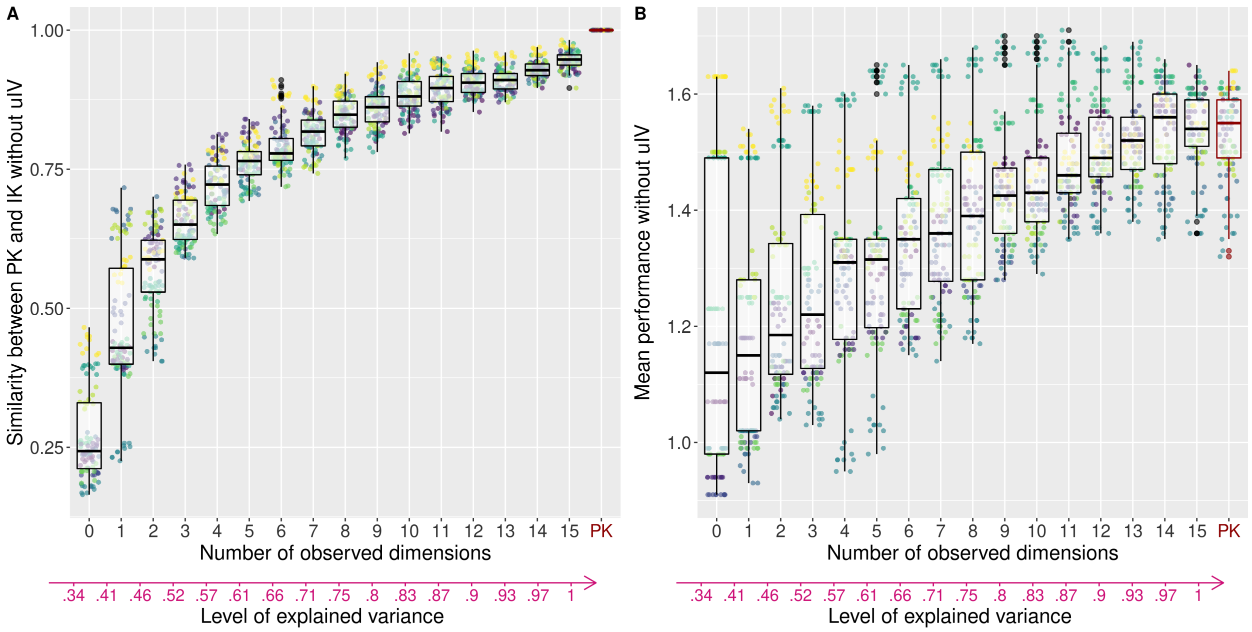


**Figure S4.16: Effect of the number of observed environmental dimensions on the similarity in final species abundances between models and on the site sorting.** Each color represents an $E \times S$ configuration. For the similarity (A), each point represents the pairwise percentage similarity (PS) in the final species abundances between two simulations with the same $E \times S$ configuration and the same initial conditions (ten points per color), but obtained using the *Perfect knowledge model* one the one hand and one of the *Imperfect knowledge models without uIV* on the other hand. For the site sorting - right panels -, each point represents the community mean performance of the final communities. This mean performance was calculated with the *Perfect knowledge model* and averaged across all individuals at the end of the simulation. These two metrics were computed for communities simulated with the *Imperfect knowledge models without uIV* (0 to 15 observed dimensions) and with the *Perfect knowledge model* (PK, red, far right). Results shown here were obtained with a *deterministic mortality* and an *abundance-dependent fecundity*.

### 5 Spatial illustration of the experiment

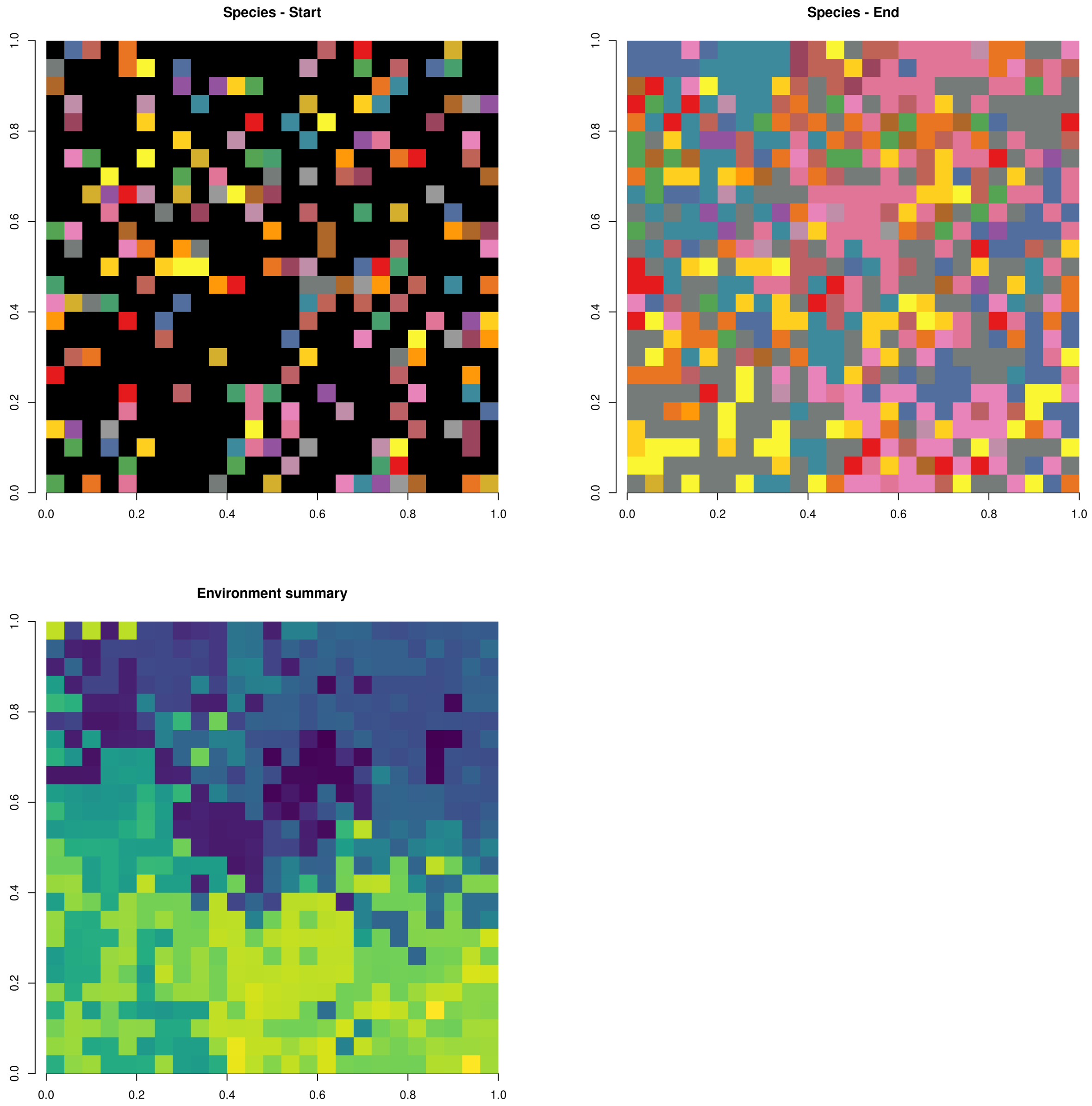


**Figure S5.17: Example of the spatial distribution of species at the beginning (top left) and at the end (top right) of a simulation, and of the environment (bottom).** In all panels, each square is a site. In the top panels, each colour represents a species, except black which represents empty sites. In the bottom panel, colours represent a summary variable of the fifteen environmental dimensions. This variable was computed using the three first principal components of a PCA, which were considered as RGB values and were combined in order to create a unique result for each combination while also minimising the dominance of one RGB value over the others. We thank Émeric Tourniaire for the idea and the code of the combination. Results shown here were obtained with the *Pefect knowledge model*, using a *deterministic mortality* and an *abundance-dependent fecundity*.
